# Lipopolysaccharide Precursor Mutants Disrupt Unipolar Polysaccharide Adhesin Synthesis and Cell Surface Functions in *Agrobacterium tumefaciens*

**DOI:** 10.64898/2026.06.01.729274

**Authors:** Ian P. Reynolds, Isabel M. Westin, Caleb J. Wood, Clay Fuqua

## Abstract

*Agrobacterium tumefaciens* is a facultative phytopathogen and causative agent of crown gall disease, capable of attaching to surfaces via a polarly localized unipolar polysaccharide (UPP) adhesin. The UPP is composed of two distinct forms, one characterized by *N*-acetylglucosamine residues (UPP_GlcN_) and the other by *N*-acetylgalactosamine residues (UPP_GalN_). Two genes with presumptive roles in dTDP-L-rhamnose biosynthesis, *rfbA* and *rfbD,* were identified in a transposon screen to be involved specifically in UPP_GalN_ biosynthesis. Examination of independent *rfbA* and *rfbD* mutations validates the transposon mutant phenotypes, with UPP_GalN_ specific defects observed as well as general attachment defects and UPP-mediated cellular aggregation across all *rfb* mutants. Additionally, despite retaining flagella, mutations in the *rfb* genes impart flagellar motility defects. Suppressor mutations that rescue the non-motile Δ*rfbD* phenotype disrupt the remaining *rfb* genes (*rfbA*, *rfbB,* and *rfbC*), the phosphoglucomutase *exoC* and a putative glycosyl transferase ATU-RS21610. Single deletion mutants in the dTDP-L-rhamnose pathway also have hallmarks of compromised outer membrane integrity, such as increased sensitivity to high-molecular weight antibiotics and increased expression of target genes for the widely conserved ChvG-ChvI two component system outer membrane stress response. Thus, defects in the dTDP-L-rhamnose pathway cause multiple cell surface deficiencies, resulting in outer membrane stress, impaired flagellar motility, defective UPP production, and irregular adhesion.

**Importssssance:** Biofilms are clinically and industrially relevant in many different contexts, with substantial health impacts and monetary costs. Surface attachment and motility are essential components necessary for bacterial biofilm formation. Many microbes utilize specific surface appendages or structures to facilitate attachment and motility. This study probes the process of surface attachment and subsequent biofilm formation in *A. tumefaciens* and has revealed connections between the creation of polysaccharide precursors required for biofilm formation and outer membrane functions such as motility and membrane integrity. These findings broaden our understanding of the complex interactions between outer membrane surface functions and how disruption of relevant precursor pools can have multifaceted impacts that may represent useful targets for new antimicrobial approaches.

## Introduction

Bacterial biofilms are communities of surface-attached cellular aggregates characterized by dramatic physiological changes relative to planktonic cell growth. Biofilm formation is thought to protect bacteria from a variety of environmental stressors, such as toxins, antibiotics, and predation, and can additionally serve as a strategy for increasing nutrient acquisition (1–3). These communities often contain and are surrounded by a self-derived extracellular matrix of exopolysaccharides, extracellular DNA and protein components. The propensity for biofilms to form upon living surfaces is medically relevant, as the same mechanisms protecting these communities render them more resistant to therapeutic strategies (4–6). During host interactions, surface attachment and biofilm formation are often essential steps for colonization or pathogenesis. A common molecular mechanism for surface attachment and subsequent biofilm formation is the production of adhesins, such as pili, flagella, exported proteins and exopolysaccharides, and these structures often also mediate bacterial interactions with eukaryotic hosts (7–9).

*Agrobacterium tumefaciens* is member of the Alphaproteobacteria (APB) and is the causative agent of crown gall disease in dicot plants, with infection resulting in neoplastic growth of the infected tissue. Release of energy-rich metabolites from infected tissues, termed opines, serves as a custom nutrient source for the infecting *A. tumefaciens* populations (10–13). The *A. tumefaciens* type strain C58 (also called *A. fabrum* or genomospecies G8) encodes for production of at least five non-essential exopolysaccharides including cellulose, succinoglycan, cyclic and linear β-1-2 glucans, curdlan (β-1,3 glucan), and a polarly localized polysaccharide-based adhesin called the UPP (<u>u</u>ni<u>p</u>olar <u>p</u>olysaccharide). Biofilm formation through UPP and cellulose contribute to stable interactions with plant hosts during pathogenesis, though the UPP alone is sufficient for stable surface attachment on both biotic and abiotic surfaces (14–16). UPP-type polysaccharide genes are conserved within the *Alphaproteobacteria* (APB) phylum, with both mammalian pathogens such as *Brucella melitensis* and *Ochrobacterum anthropi,* and plant-associated microbes such as *Rhizobium leguminosarum, Bradyrhizobium japonicum*, and *Sinorhizobium meliloti*, all retaining gene clusters with homologs of UPP biosynthetic genes (15, 17, 18).

Our prior studies identified a Wzx-Wzy UPP biosynthesis pathway, driving production of two chemically, spatially, and genetically distinct exopolysaccharides (14, 19). Each UPP species is defined by binding ability to lectins specific for either *N*-acetylglucosamine (GlcNAc; WGA - wheat germ agglutinin) or *N*-acetylgalactosamine (GalNAc, DBA - *Dolichos biflorus* lectin). The GlcNAc-containing UPP (UPP_GlcN_) is polymerized by the Wzy-type polymerase UppW (ATU_RS11495), and in contrast the polymerization of the GalNAc-containing UPP (UPP_GalN_) requires the Wzy-type polymerase UppY (ATU_RS02370). UPP production is surface contact-inducible and controlled by the cytoplasmic second messenger c-di-GMP (14, 16). The UPP is overproduced in a background genetically disabled for the other non-essential exopolysaccharides (cellulose, curdlan, β-1,2 glucan, succinoglycan; CDGS-, UPP+) and elevated for c-di-GMP (due to a regulatory mutation in the Δ*pruA* gene (20)). A transposon mutant screen of the CDGS- Δ*pruA* derivative was performed on solid medium containing Congo Red (CR) dye which binds to both UPP_GlcN_ and UPP_GalN_. A <u>d</u>ecrease in <u>C</u>ongo <u>R</u>ed (DCR) staining is well correlated with decreased UPP production and an attachment deficiency (14). The UPP_GlcN_ and UPP_GalN_ species are each sufficient to impart CR staining and two parallel transposon mutant screens were performed in CDGS- Δ*pruA*Δ*uppY* and CDGS- Δ*pruA*Δ*uppW* backgrounds. These screens identified mutations that impact synthesis of both polysaccharides, as well as candidate genes for additional pathway-specific functions that independently affect either UPP_GalN_ or UPP_GlcN_ (14). Two of the genes identified in the UPP_GalN_ (Δ*uppW*) screen, were *rfbA* (ATU_RS21635) and *rfbD* (ATU_RS21640), which encode a glucose-1-phosphate thymidylyltransferase and a dTDP-4-dehydrorhamnose reductase, respectively (14). These genes are annotated to function in the production of dTDP-β-L-rhamnose, which serves as a substrate of the O-antigen and/or core oligosaccharide of lipopolysaccharide (LPS) (21). In *A. tumefaciens* C58, it is reported that rhamnose is a component of the core oligosaccharide, with a linear homopolysaccharide of 3-α-L-6dTalose serving as the O-antigen (22).

In bacteria, rhamnose is frequently utilized in separate biological processes, such as biosurfactant production, outer membrane biogenesis, and exopolysaccharide biosynthesis, and this broad utilization can lead to pleiotropic effects when perturbing the *rfb* genes (23–26). Indeed, the impact of disrupting dTDP-β-L-rhamnose synthesis has been explored in bacteria such as *Salmonella enterica, Pseudomonas aeruginosa,* and the APB *Caulobacter crescentus,* and disruption of *rfb* genes can consequently impact processes such as LPS/LOS biogenesis, surface attachment and flagellar motility (27–30). Considering the role of the *rfb* genes in sugar precursor biosynthesis and more broadly potential impacts on cellular processes such as attachment and motility, it seemed plausible for these genes to function either directly or indirectly within UPP precursor pools, with a more pronounced effect on the UPP_GalN_ specific precursor pool. We sought to explore the specific function of the *rfb* genes, in particular the impact of the *rfbA and rfbD* genes on synthesis of UPP_GalN_ relative to that of UPP_GlcN_, and found that these genes affect both UPP-dependent surface attachment and more broadly outer membrane surface functions including flagellar motility and antibiotic exclusion in *A. tumefaciens*.

## Results

### The *A. tumefaciens rfbCBDA* gene cluster and the presumptive dTDP-β-L-rhamnose biosynthetic pathway

In the *A. tumefaciens* C58 genome the *rfbA* and *rfbD* genes are encoded within the four-member operon *rfbCBDA* (Fig. 1A) along with *rfbC* (ATU_RS21650) and *rfbB* (ATU_RS21645). RfbC is predicted to be a dTDP-4-dehydro-6-deoxy-glucose-3,5-epimerase and RfbB is a dTDP-glucose 4,6-dehydratase, each driving different steps in dTDP- β-L-rhamnose synthesis. The coding sequences in the *rfbCBDA* operon are very tightly linked and likely to be translationally coupled in *A. tumefaciens*. The *rfb* genes are well conserved among Proteobacteria including the APB, although the gene order can vary (31). Biosynthesis of dTDP-β-L-rhamnose is thought to proceed from dTTP and glucose-1-phosphate via reactions catalyzed in sequence by RfbA, RfbB, RfbC and RfbD (Fig. 1B) to generate the sugar nucleotide precursor for addition of rhamnose residues to LPS/LOS. Thus, inactivation of *rfbA* would block the first committed step of the pathway producing dTDP-glucose, and *rfbD* would block the final step reducing dTDP-4-keto-L-rhamnose to dTDP-L-rhamnose. However, because of the *rfbCBDA* operon structure, transposon insertions in *rfbD* are likely to be polar on expression of *rfbA*, and thus the *rfbD* transposon mutant phenotypes could reflect impacts on both genes.

**Figure 1:**
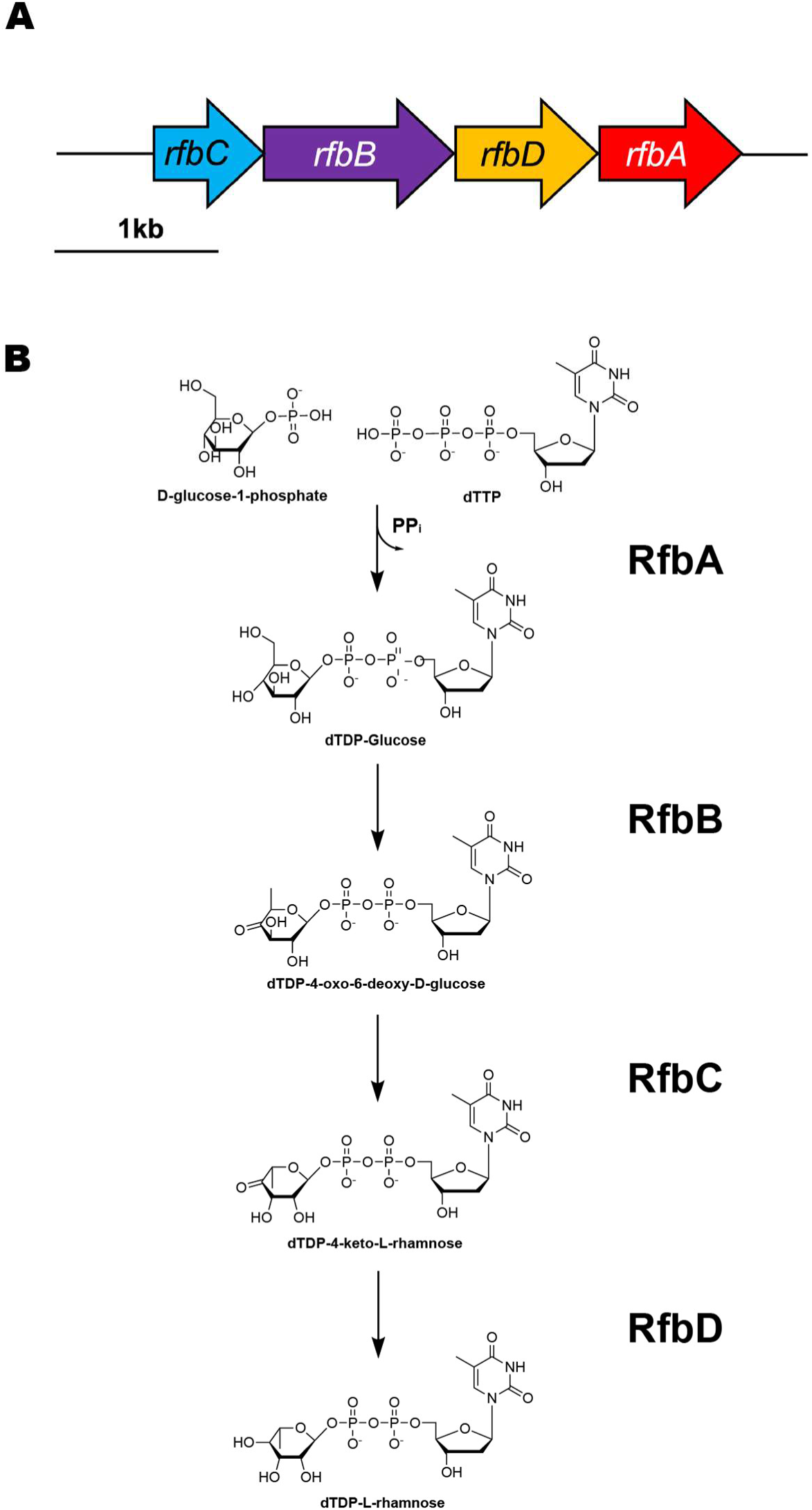
Biosynthetic pathway and operon structure of *rfb* genes. A) The operon structure of the *rfbCBDA* operon. B) The biosynthetic pathway beginning from the initial addition of dTTP to G-1P by RfbA through to the production of dTDP-B-L-rhamnose by RfbD. Adapted from Raetz (21).

### The *rfbCBDA* operon contributes to polysaccharide-dependent phenotypes in *A. tumefaciens*

While *rfbD* and *rfbA* were two genes identified as putative UPP determinants in our screen, the full *rfbCBDA* operon was examined for surface attachment and CR binding as a proxy for UPP production. The CR staining of each *rfb* mutant was measured against a deletion mutant of the entire operon (Δ*rfbCBDA*) and wild-type *A. tumefaciens* C58, in addition to *rfb* mutants within the elevated c-di-GMP background of C58 CDGS- Δ*pruA*. The *rfb* mutants in wild type in which the UPP and cellulose are typically produced at low levels show slightly increased CR staining, suggesting feedback into other CR-labeled polysaccharide biosynthesis pathways such as cellulose (Fig. 2A). Loss of *rfbC* confers the greatest increase in staining, with the most modest increases being in Δ*rfbA* and Δ*rfbD*. Ectopic expression of each corresponding gene (or expression of the full operon in the Δ*rfbCBDA* mutant) rescues CR phenotypes. Additionally, these *rfb* mutants generally have visible pellicle formation, indicative of increased cellulose production, as observed in the sheet-like growth at the air-liquid interface in the Δ*rfbA,* Δ*rfbB* and Δ*rfbCBDA* mutants (Fig. 2B). In the C58 CDGS- Δ*pruA* mutant, deletion of any *rfb* genes appears to modestly decrease CR staining. The greatest decreased staining is shown in the C58 CDGS- Δ*pruA* Δ*rfbD* mutant, and complementation likewise restores CR staining to that of the parent strain (Fig. 2D). Ectopic expression of complementation plasmids also rescues these phenotypes in the other mutants. While decreased CR staining suggests less UPP is present, these *rfb* mutants display notable cellular aggregation in liquid media, with each mutant markedly increased relative to the parent strain (Fig. 2E).

**Figure 2:**
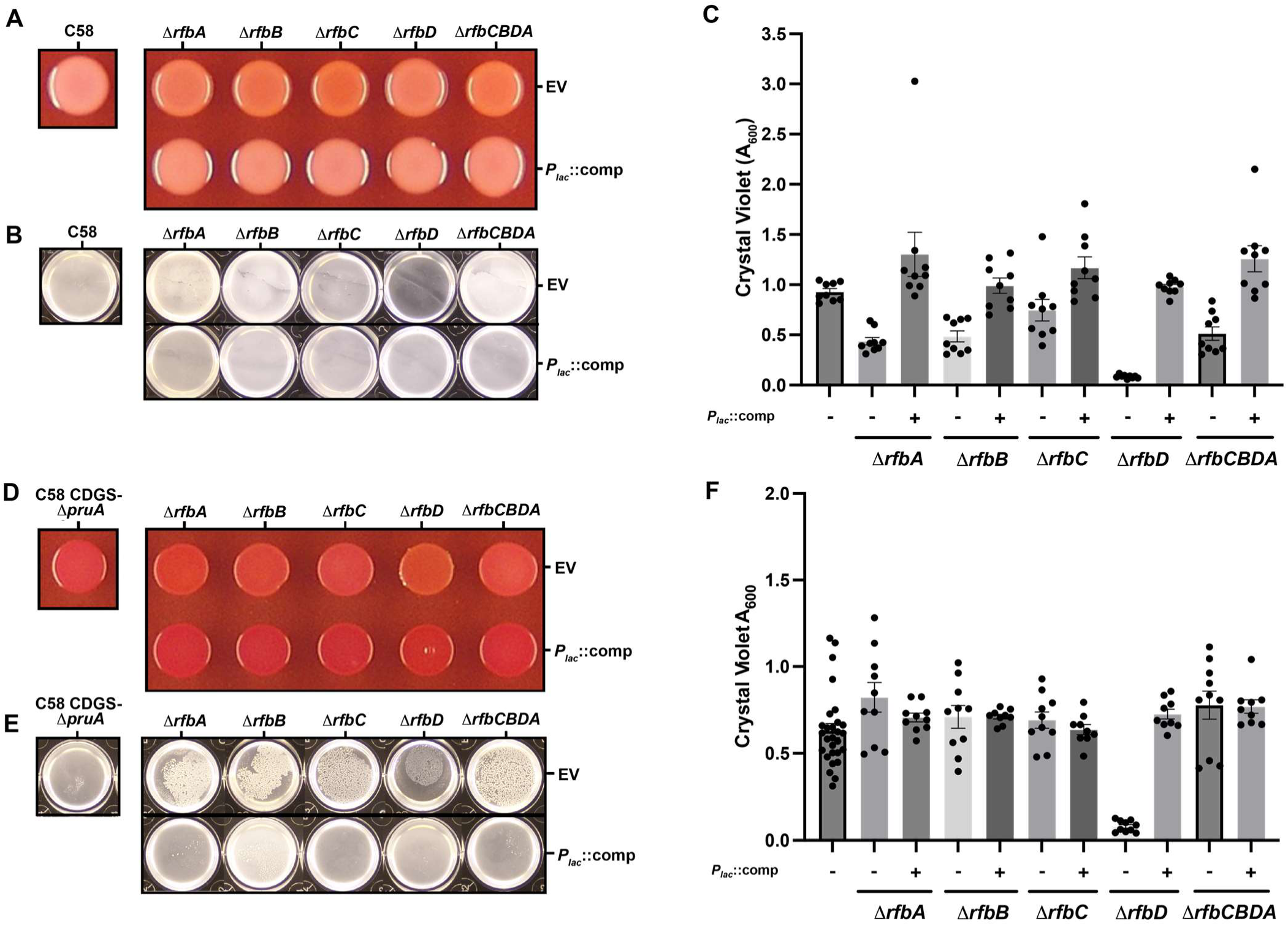
The *rfbCBDA* operon contributes to polysaccharide production, pellicle growth, and biofilm formation. (A) The Congo Red (CR) staining phenotypes of WT C58 *rfb* mutants plated on ATGN-CR (75 ug/mL). Top row: Colonies with the pSRKGm vector (EV) Bottom row: Corresponding *rfb* gene complement and vector controls are ectopically expressed with 500 *µ*M IPTG. (B) Representative images of pellicle formation observed during biofilm formation at 48 h are shown below, designated by the same schema as in A. C) Biofilm formation on PVC coverslips after 48 h incubation for WT C58 with each *rfbCBDA* gene deleted, and the full operon deletion. Adhered cells were stained in Crystal Violet (CV), solubilized in 30% acetic acid, with A_600_ measured. Plasmid-borne copies of *rfb* genes under *P_lac_* control induced with 500 *µ*M IPTG. D) CR staining phenotypes of *rfb* mutations in C58 CDGS- Δ*pruA* backgrounds, as described for panel A. E) Representative images of pellicle formation during biofilm formation assays at 48 h as in panel B. F) Biofilm formation of *rfb* mutants in C58 CDGS- Δ*pruA* backgrounds. Adhered cells on PVC coverslips were stained with CV as described in panel A. For panels C and F, bars are means of biological triplicate assays, and the error bars are SEM. Data for the parent strain (C58 CDGS- Δ*pruA*) biofilm assay experiments was compiled from a larger numbers of repeats.

Biofilm formation assays using crystal violet (CV) staining of attached biomass to polyvinyl chloride coverslips were performed with *rfb* mutants in both wild-type C58 and in the CDGS- Δ*pruA* mutant. Loss of *rfbA* and *rfbD* in wild-type C58 decreases biofilm formation (p-values of 0.02 and <0.0001, respectively, one-way ANOVA), while *rfbB*, *rfbC*, *and rfbCBDA* show statistically non-significant decreases (p value > 0.05 for all, one-way ANOVA) (Fig. 2C). When examining *rfb* mutants in C58 CDGS- Δ*pruA* backgrounds the Δ*rfbD* mutant is dramatically decreased (p value <0.0001, one-way ANOVA), but despite the clear increased auto-aggregation and pellicle formation in the other *rfb* mutants, there are no significant changes in surface attachment (Fig. 2F). While CR staining generally correlates well with UPP production and the downstream impact on biofilm formation (14), only in the C58 CDGS-Δ*pruA* Δ*rfbD* mutant that is most decreased for CR staining do we observe significantly impaired surface attachment.

### Polysaccharide staining phenotypes in *rfbA* and *rfbD* UPP-polymerase mutants

The *rfbA* and *rfbD* genes were initially identified due to their DCR mutant phenotypes in the CDGS- Δ*pruA* mutant in which the UppW polymerase was also disabled (14). To further examine the extent to which *rfbA* and *rfbD* contribute to UPP_GalN_ or UPP_GlcN_ phenotypes, in-frame deletion mutants were generated in mutants for the UPP Wzy-polysaccharide polymerases (Δ*uppY,* Δ*uppW,* and the Δ*uppY* Δ*uppW* double mutant) for both wild type C58 and the CDGS- Δ*pruA* mutant. Indeed, the decreased CR (DCR) phenotype of Δ*rfbA* and Δ*rfbD* mutants in the CDGS- Δ*uppW*Δ*pruA* mutant validates their isolation from the transposon mutant screen, with the Δ*rfbA* mutant phenotype modestly DCR and Δ*rfbD* mutant with a more pronounced phenotype (Fig. 3A). Aside from the expected DCR phenotype in Δ*uppW*Δ*pruA* CDGS-, there appears to be general decreased staining in the Δ*uppY*Δ*pruA* CDGS-. The residual CR staining in these mutants is however completely abolished in the Δ*uppY*Δ*uppW* double mutant. The UPP-specific phenotypes observed in these polysaccharide-deficient mutants show the effect of *rfb* mutants independent of their impact on other polysaccharide pathways. As seen in the single gene knockouts of *rfb* genes in wild-type C58, loss of *rfbA* and *rfbD* in UPP-polymerase mutant backgrounds retaining EPS production generally caused a slight increase in CR staining (Fig. S1A) with the lone exception being a minimal change in CR staining in the Δ*uppY*Δ*uppW*Δ*rfbD* mutant. The increased staining of *rfb* mutants in otherwise wild-type backgrounds is distinct from the decreased CR staining observed in the Δ*pruA* CDGS-backgrounds in which the only non-essential characterized polysaccharide substantially produced is UPP.

**Figure 3:**
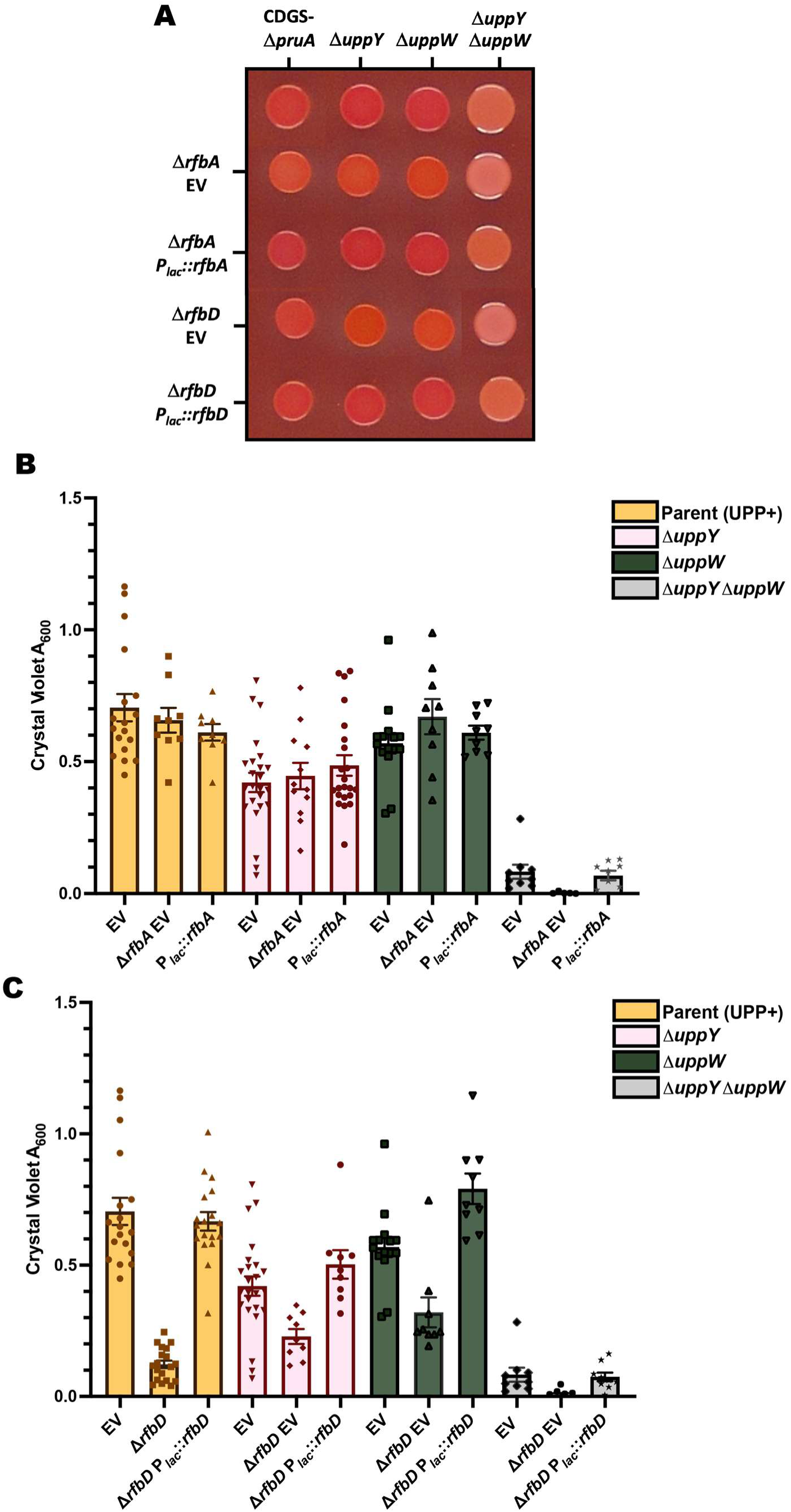
Attachment and Congo Red staining of *rfbA* and *rfbD* mutations in UPP polymerase mutants. (A) CR staining of each *rfbA* and *rfbD* mutant in the C58 CDGS-Δ*pruA* mutant with Wzy-type polymerase mutations. Cultures were spotted onto ATGN-CR (75 *µ*g/mL) plates with 500 µM IPTG for 48 h. B) Biofilm formation on PVC coverslips after 48 h incubation for Δ*rfbA* mutant in UPP Wzy-polymerase mutants in the C58 Δ*pruA* CDGS- screening strain (elevated c-di-GMP and EPS-). CV staining of adhered cells was solubilized in 30% acetic acid and the A_600_ was measured for parent strains and *rfbA* mutants in C58 CDGS- Δ*pruA* genetic backgrounds with Δ*uppY*, Δ*uppW*, and Δ*uppY*Δ*uppW* Wzy-type polymerase deletions. Plasmid-borne copies of *rfbA* were under the control of *P_lac_* and induced with 500 *µ*M IPTG. EV, pSRKGm control. (C) CV staining of 48 h PVC coverslip biofilms for Δ*rfbD* mutant in UPP Wzy-polymerase mutants in the C58 Δ*pruA* CDGS- screening strain. Performed as described in panel B. For panels B and C, bars are the mean of biological triplicate assays, and the error bars are SEM.

### Mutations in *rfbA* and *rfbD* have differential impacts on UPP-mediated biofilm formation

We hypothesized that loss of either of the *rfbA* and *rfbD* genes would manifest in decreased attachment and biofilm formation given the isolation of these genes and the generally reliable correlation between CR staining binding and attachment phenotypes in our prior studies (14, 15). Biofilm assays were performed in the same UPP-polymerase mutants described above to evaluate any effects on surface attachment for the Δ*rfbA* and Δ*rfbD* mutants. Surprisingly, the Δ*rfbA* mutation minimally impacts attachment in the Δ*pruA* CDGS- mutant, with no significant effect of the additional *uppY and uppW* mutations, though the Δ*uppY*Δ*uppW* double mutant remains incapable of attachment (Fig. 3B). In contrast, loss of *rfbD* in the elevated c-di-GMP background leads to a statistically significant decrease in attachment in the Δ*uppW* mutant and otherwise wild type backgrounds (p-value <0.05, one way ANOVA) (Fig. 3C). There is a significant decrease for the Δ*uppY*Δ*rfbD* mutant (p value 0.015, one-way ANOVA) and no significant difference within the Δ*uppY*Δ*uppW* mutant (Fig. 3C). The observed Δ*uppW*Δ*rfbD* attachment defect is consistent with the identification of *rfbD* DCR mutants in the CDGS-Δ*pruA*Δ*uppW* screening background, suggesting a defect in UPP_GalN_ mediated surface attachment due to loss of *rfbD.* Analysis of both *rfbA* and *rfbD* mutants in the wild type shows significant attachment defects in not only the expected UPP_GalN_ only (Δ*uppW*) strains but in all UPP backgrounds, with complete loss of attachment in the Δ*uppY*Δ*uppW* double mutant (Fig. S1B, S1C). Loss of *rfbA* or *rfbD* appears to have no impact on attachment in the UPP-null mutants, confirming that attachment is UPP-dependent. While the general defects of *rfbD* knock-outs in attachment across WT and elevated UPP-producing mutants suggests a broad impact, the effect of inactivating *rfbA* appears more context specific. It may be that the auto-aggregatory phenotypes of these mutants somewhat compensate for the UPP deficiencies revealed in the Wzy-polymerase mutants.

### Stimulation of UPP-mediated aggregation

Coinciding with impacts on biofilm formation, loss of either of the *rfbA* and *rfbD* genes drastically increases auto-aggregation and pellicle formation in liquid culture. This increased aggregation is lost in the Δ*uppY*Δ*uppW* double mutant. The Δ*uppW* mutant without *rfbA* or *rfbD* disruption also displays a higher baseline aggregation, that is dramatically increased by the Δ*rfbA* and Δ*rfbD* mutations (Fig. 4A). Interestingly, the Δ*rfbD* aggregation is abolished by the Δ*uppY* mutation whereas Δ*rfbA* aggregates with either UPP species present. Aggregation due to *rfbA* and *rfbD* mutation is most dramatic in the elevated c-di-GMP screening background, but diminished or absent in the wild type.

**Figure 4:**
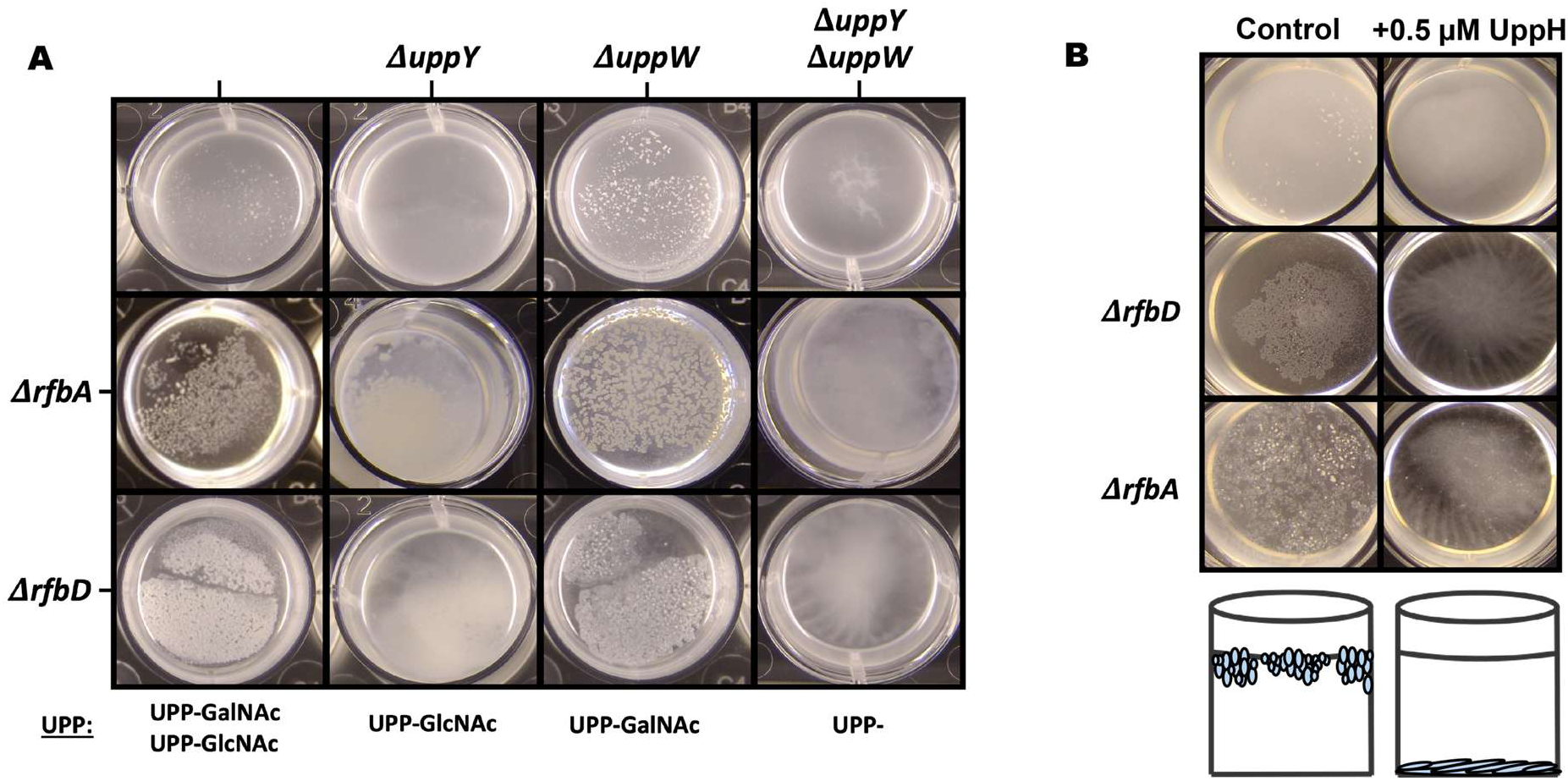
UPP Mediated aggregation and pellicle formation in *rfb* Mutants. (A) Pellicle formation in C58 CDGS- Δ*pruA* strains with Δ*uppY*, Δ*uppW*, and Δ*uppY*Δ*uppW* Wzy-type polymerase deletions. Samples were incubated in ATGN at RT for 48 h prior to imaging. Columns and rows labeled with Wzy polymerase and *rfb* gene mutations, respectively. (B) Pellicles formed by C58 CDGS- Δ*pruA* derivatives (*rfb* genotypes as indicated) incubated with purified UppH glycoside hydrolase (0.5 μM). Samples were treated with UppH enzyme or the equal volume of buffer (control column) for 48 h at RT prior to imaging. Schematic of pellicle or sedimented cells is shown.

It is plausible that the auto-aggregation observed in *rfb* mutants is due to disruptions in LPS precursor biogenesis and changes in outer membrane composition, rather than directly due to the UPP adhesin (29). Although Δ*rfbA and* Δ*rfbD* mutants fail to aggregate in the UPP-Δ*uppY*Δ*uppW* double mutant, combinatorial synthetic effects due to multiple mutations are also possible. As an additional test, we incubated these cultures in the presence of purified UppH, a glycoside hydrolase that can degrade the UPP when provided exogenously (14). Co-incubation with purified UppH resolves the aggregatory phenotype of *rfbD* and *rfbA* mutants, including for Δ*uppW* mutants where aggregation is most severe (Fig. 4B). Significant sedimented biomass remains visible in the wells of cell culture dishes but the pellicle is fully dispersed.

### Impaired motility in *rfbCBDA* mutants

The notable sedimentation of UppH-treated cultures for the *rfb* mutants relative to wild type *A. tumefaciens* (Fig. 4B) hinted at defects in flagellar motility. It has previously been reported that loss of *rfb* genes in other bacteria can decrease motility, although the mechanism by which this occurs is unclear (32–34). Indeed, swim assays on 0.25% motility agar revealed that the Δ*rfbD* mutant is non-motile, whereas the Δ*rfbA* mutant is decreased relative to wild type but retains weak motility (Fig. 5A). Additionally, mutants in the remaining *rfb* operon genes, *rfbB*, *rfbC*, and the full operon *rfbCBDA*, display defective motility. Motility appears graded across *rfb* mutants, with loss of *rfbC* and *rfbD* being most deleterious while Δ*rfbA* and Δ*rfbB* mutants display more modest defects. Deletion of the entire *rfbCBD*A operon (Δ*rfbCBDA*) itself results in weak motility (Fig. 5A). One explanation for the variability of motility deficiency in the different *rfb* mutants, would be partial rescue by paralogous *A. tumefaciens* genes. There are close paralogs to *rfbA* (ATU_RS18920) and *rfbB* (ATU_RS21620) in the *A. tumefaciens* C58 genome that could act in the absence of the native gene, moderating the mutant phenotypes. In contrast, there are no close paralogs to *rfbD* encoded in the genome. Compensation appears to be the case for *rfbA* with ATU_RS18920 (*exoN)* as deletion of this paralog further worsens the motility defect in the Δ*rfbA* mutant (Fig. S2). However, the Δ*rfbA*Δ*exoN* double mutant still displays stronger motility than the Δ*rfbD* mutant. Testing for other paralogous pairs such as *rfbB* with ATU_RS21620, failed to reveal such additive effects (Fig. S2).

**Figure 5.**
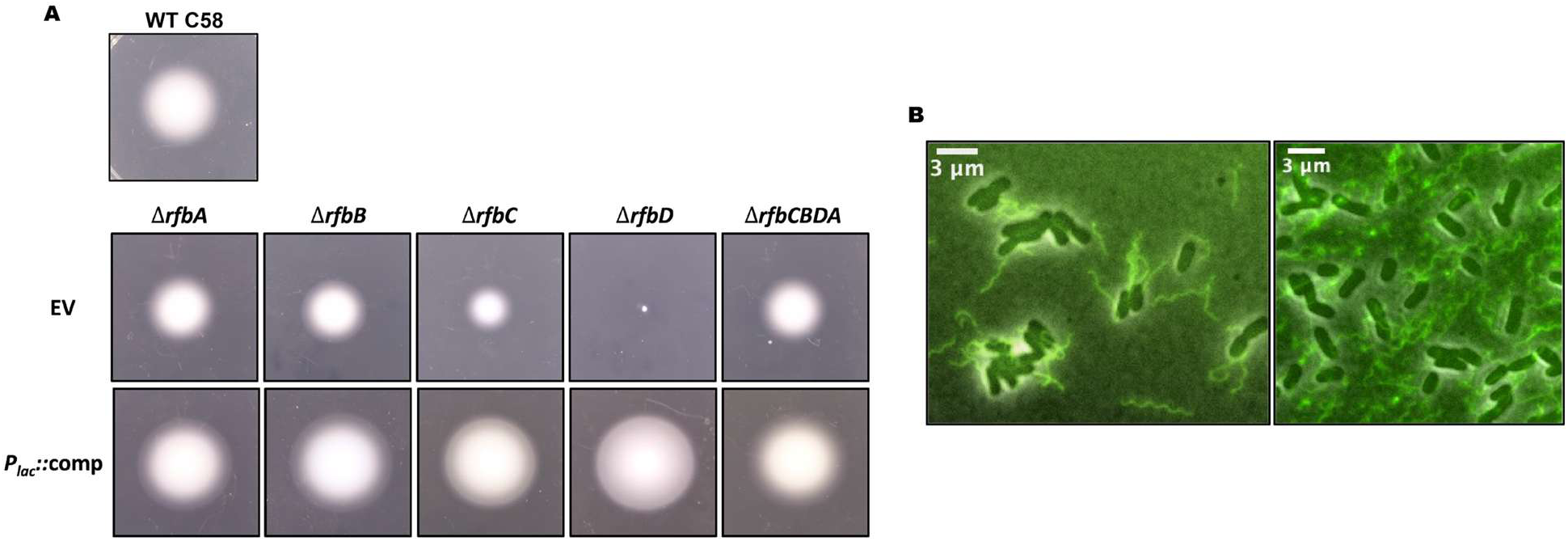
Defective flagellar-mediated motility in *rfb* mutants. (A) Representative motility assays of *rfb* mutants. A single colony for each sample was inoculated into 0.25% agar ATGN and incubated at RT for 3 days prior to imaging *P_lac_*::comp indicates each respective pSRKGm complementation construct, induced with 500 *µ*M IPTG. EV, empty pSRKGm vector. (B) Flagella visualization in C58 Δ*rfbD* mutants (left panel) and in WT FlaAT213C (right panel) strains harboring pBM205 (pSRK::*P_lac_*:: FlaAT213C + *P_flaA_* native promoter) labelled with 0.5 *µ*g/mL AF-488 maleimide as described in Materials and Methods. Each image was captured with a Nikon E800 epifluorescence microscope under 100x oil immersion objective.

Observation of planktonic cultures under the microscope revealed no visible motility for the Δ*rfbD* mutant and sporadic motility for the Δ*rfbA* mutant. These defects were fully complemented by provision of the wild type *rfbA* and *rfbD* genes. Despite these motility deficiencies, even in the case of the severely deficient Δ*rfbD* mutant, these cells retain flagellar biosynthesis as viewed by phase contrast microscopy of a phenol-based flagellar stain and by fluorescence microcopy of a Cysteine knock-in flagellin (*flaA*-Cys) labeled with a cysteine reactive dye (Fig. 5B) (35). There is a clear connection between motility and surface adherence, and powered flagella are necessary for biofilm formation in *A. tumefaciens* under static conditions (36), and robust biofilm formation in other biofilm models such as *E. coli* and *Pseudomonas* spp. also requires active motility (37, 38). Diminished flagellar motility could therefore impart decreased biofilm formation, though the capacity for surface adherence upon reaching a surface is less affected, as an aflagellate *A. tumefaciens* mutant under flowing conditions forms robust biofilms (36). The relationship between the motility and adhesin production phenotypes demonstrated by the *rfb* mutants could both impact subsequent biofilm formation.

We sought to determine whether the motility defects could be related to the aberrant UPP production in these mutants, such as through UPP-dependent aggregation. In all backgrounds tested, *rfbA* mutants displayed UPP-independent motility defects within wild type backgrounds disabled for the various UPP-species (Fig. S3A-B). However in otherwise wild-type C58 Δ*rfbD* when *uppW* is absent, such as the Δ*uppW* and Δ*uppY* Δ*uppW* mutants, the motility defect is suppressed (Fig. S3C, S3D). The reason for this suppression remains difficult to explain and the isolation of Δ*rfbD* motile suppressor mutants with unbiased transposon mutagenesis did not identify *uppW* mutants (see below). Examining these mutants within CDGS- Δ*pruA* backgrounds suggests that the effects of *rfb* mutations on motility are distinct from the effects on UPP biosynthesis, and changes in c-di-GMP levels between the wild type and the *pruA* CGDS- strain are also excluded (Fig. S4). Motility signatures are otherwise consistent within the *rfbA* mutants (Fig. S4A-B), with partial motility defects, whereas Δ*rfbD* mutants have more severe losses (Fig. S4C-D), in each case regardless of UPP-species present. C58 CDGS- Δ*pruA* strains overall appear to have modestly attenuated motility relative to WT and mutants in *rfb* genes in these backgrounds therefore display a lower motility baseline (likely impacted by the elevated c-di-GMP and absence of cyclic and linear β-1,2 glucans in these mutants; Fig. S4A-D).

### Enrichment for motility suppressors implicates dTDP-β-L-rhamnose metabolism

We sought to investigate the mechanistic basis of the *rfbCBDA* mutants using a genetic suppression analysis taking advantage of the dramatic motility defect in the Δ*rfbD* mutant.The C58 CDGS- Δ*pruA*Δ*rfbD* and C58 Δ*rfbD* mutants were subjected to transposon mutagenesis and used to generate *Mariner* transposon mutant libraries (>21,000 colonies). A homogeneous suspension of each library was plated on 0.25% ATGN motility agar and incubated 48 h at RT to allow for enrichment of motile suppressors within the population. This extended incubation allowed for enrichment of transposon mutants that suppressed the non-motile phenotype of this mutant, appearing as swim rings that emerged slowly (Fig. 6). Incubation of the C58 CDGS-Δ*pruA* Δ*rfbD* strain on this medium also fostered formation of visually striking tendrils that extended from the swim ring, but these were distinct from the swim rings formed by true suppressor mutants and appear to be a flagella-dependent feature of plating dense culture volumes of this UPP-overproducing genotype (Fig. S5).

**Figure 6:**
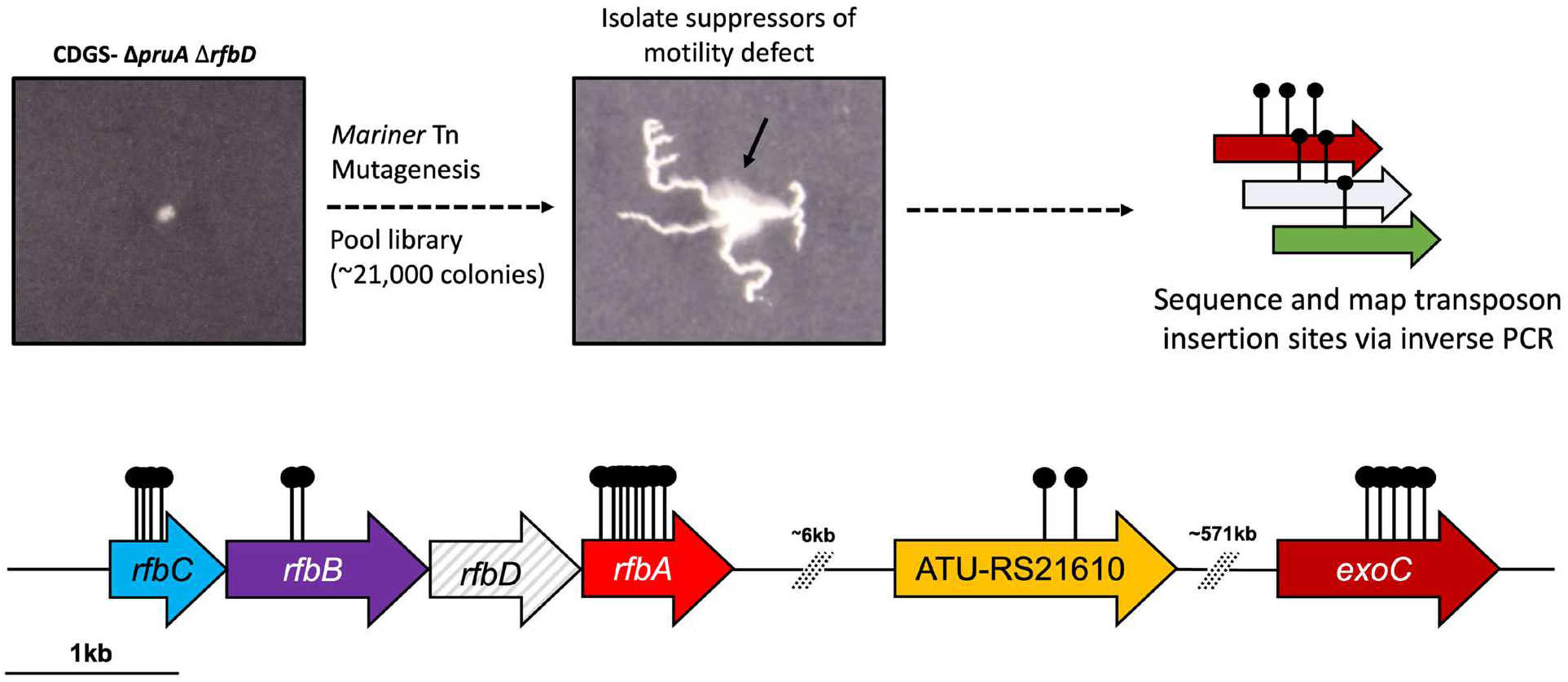
Transposon mutagenesis and enrichment for motility suppressors in the nonmotile CDGS-Δ*pruA* Δ*rfbD* mutant. A *Himar1* transposon library was prepared in the non-motile C58 CDGS- Δ*pruA* Δ*rfbD* mutant and 5 μL of library was incubated on 0.25% ATGN motility agar ATGN for 48 h at RT prior to sampling from the late forming motile swim ring for suppressors of the motility defect. Transposon insertions in these motile derivatives were mapped via inverse PCR as described in Materials and Methods.

Motility suppressors were collected from the nascent swim ring, isolated, and verified for motility by inoculating onto fresh 0.25% ATGN motility agar. Genomic DNA was then isolated, and transposon insertion sites were mapped by inverse PCR as described in the experimental methods. Strikingly, the primary means for suppression of the severe motility defect in the C58 CDGS- Δ*pruA*Δ*rfbD* mutant is disruption of the remaining *rfb* genes (*rfbC, rfbB, rfbA*). Eight independent insertions were identified within *rfbA*, the initial step within the biosynthetic pathway (Table S1). Multiple transposon insertions into *rfbC* and *rfbB* were also observed, suggesting that interruption of any of the preceding enzymatic reactions can suppress the defect imposed by loss of *rfbD.* Five independent transposon mutants were isolated in *exoC* (ATU_RS19035), the phosphoglucomutase that converts glucose-6-phosphate into the common precursor glucose-1-phosphate that is utilized as a substrate for *rfbA* (Fig. 1B). Nonpolar deletion of *exoC* itself within wild type C58 results in pleotropic defects, including decreased production of succinoglycan, a motility defect, and deficiencies in UPP production (14). Two independent insertions mapped to the GT2 domain of a hypothetical glycosyl transferase (ATU_RS21610, Atu4610) encoded 6354 bp away from the *rfbCBDA* operon. While not obviously involved directly in metabolism of the Rfb pathway, this protein is predicted to have dual-domains, a methyltransferase and a GT2, and has sequence homology to genes annotated to function in cell wall biogenesis. Preliminary analysis of transposon mutant suppressors in the otherwise WT C58 Δ*rfbD* mutant library supports the findings of the C58 CDGS- Δ*pruA*Δ*rfbD* background, with transposon insertions in the remaining *rfb* genes proving to be the primary means for suppression of the motility defect (Table S1). The only other gene with multiple insertions in this background is that of the alcohol dehydrogenase annotated as *adhP* (ATU_RS03065). Interestingly, there were no suppressor mutants identified with disruptions in *uppW* or any other UPP related genes in either transposon mutant library.

### Dysregulation of dTDP-β-L-rhamnose pathway disrupts motility and biofilm formation

Transposon insertion mutants in *rfbC and rfbB* in the remaining *rfbCBA* operon of the Δ*rfbD* parent are potentially polar on the downstream *rfb* genes. In-frame deletion mutants were generated in combination with Δ*rfbD* and each candidate partner gene (*exoC, atu4608, rfbC, rfbB, rfbA)* in the CDGS- Δ*pruA* background to validate hits from this screen. Motility defect suppression is highest in the Δ*rfbA*Δ*rfbD* mutant, with Δ*rfbB*Δ*rfbD* and Δ*rfbC*Δ*rfbD* suppressors appearing more modest relative to wild type motility (Fig 7A-B). The Δ*exoC*Δ*rfbD* mutant likewise displays visible suppression, while the ΔATU_RS21610Δ*rfbD* mutant is more subtle. The Δ*rfbA*Δ*rfbD* mutant displays motility more akin to wild type than even the full Δ*rfbCBDA* operon mutant. From this work, we see that combinatorial deletion of Δ*rfbD* with *rfbA, rfbB,* or *rfbC* can suppress the severe motility defect, but not to levels observed for the motile parent strain.

**Figure 7:**
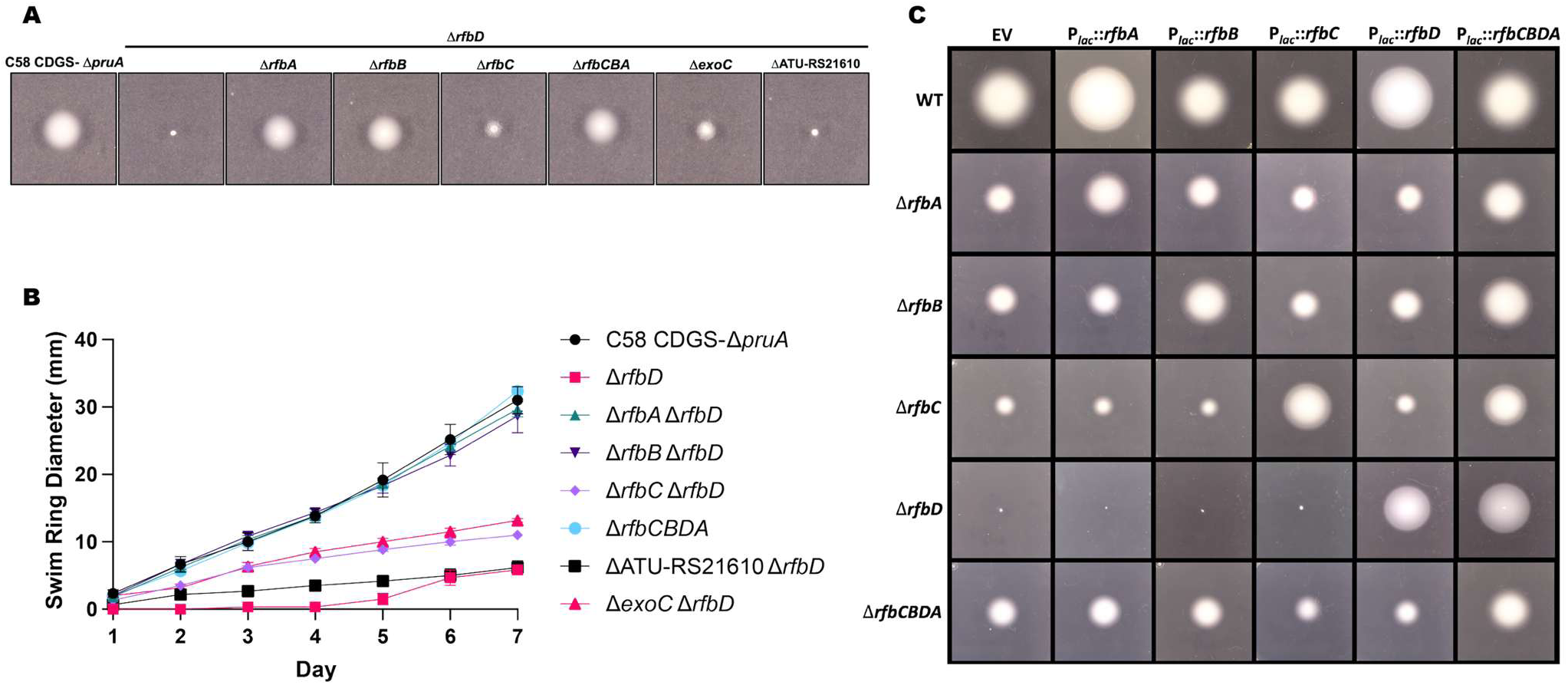
Ectopic expression of *rfb* genes and graded suppression of motility defects within paired *rfb* mutants. (A) Representative images of motility assays on ATGN agar (0.25%) of C58 CDGS- Δ*pruA*Δ*rfbD* derivatives with additional *rfbA, rfbB,* and *rfbC* deletions, relative to the parent strain and *rfbCBDA* deletion mutants. A single colony for each sample was inoculated into 0.25% agar ATGN and incubated at RT for 3 days prior to imaging. (B) Seven-day motility assay swim ring diameter values. Symbols represent the mean values performed in triplicate. (C) Ectopic expression of each *rfb* gene in each WT C58 *rfb* mutant background. Representative images captured 72 h post incubation at RT. Samples were grown as in panel B. Plasmid-borne copies of each *rfb* gene on pSRK derivates were under control of *P_lac_* and expression was induced with 500 *µ*M IPTG.

Our enrichment for motility suppressors heavily implicated the remaining *rfb* genes and the intermediate metabolites of the Rfb biosynthetic pathway. Therefore, we sought to further interrogate the relationship between motility and dTDP-β-L-rhamnose biosynthetic pathway. Ectopic expression of each *rfb* gene was performed within individual *rfb* deletion mutants as well as the full operon deletion Δ*rfbCBDA*. If the accumulation of an intermediary substrate or unregulated activity of one of these enzymes were contributing to motility defects, it was expected that ectopic expression of each gene across mutants within this pathway might correct these deficiencies. Expression constructs for each gene, and for the entire operon were introduced into the different mutants and motility was measured under inducing conditions over time. It is evident that the most severe impacts are due to loss of *rfbD* and *rfbC*, but in all cases disruption of the *rfb* pathway impacts normal motility patterns (Fig. 7C). Interestingly, single gene *rfb* operon mutants only show suppression of their motility defect with their own complementation construct, and with the complete *rfb* operon construct. With stimulation of this pathway towards production of dTDP-β-L-rhamnose by ectopically expressing *rfbCBA* or *rfbAD* within the Δ*rfbCBDA* operon mutant, we observe decreased motility even to levels similar to the nonmotile Δ*rfbD* mutant (Fig. S6).

The motile suppressors of Δ*rfbD* were also tested for their impact on the biofilm formation deficiency manifested by this mutant in otherwise wild type C58. Deletion of *rfbA* fully restored biofilm formation, and *rfbB* and *rfbC* deletions were partially restored (Fig. S7). As described above, deletion of the entire *rfbCBDA* operon partially impairs motility and likewise shows partial rescue of biofilm formation, as does deletion of ATU_RS21610. In contrast, the Δ*exoC* deletion in combination with the Δ*rfbD* mutation did not improve biofilm formation consistent with prior studies on *exoC* alone (14) even though it partially restored motility.

### An outer membrane stress response pathway is activated in *rfb* mutants

The *rfbCBDA* operon genes function in the production of an LPS precursor in many other bacteria, so we hypothesized that mutations in this pathway could result in altered outer membrane composition and in turn, outer membrane stress signaling. To test for overt changes in outer membrane composition, LOS was extracted from the WT, Δ*rfbA*, Δ*rfbD* and Δ*rfbCBDA* mutants and samples were run on an SDS-PAGE gel and silver stained to gauge for changes in LOS composition. There are no visible changes in the LOS composition between mutants, with consistent banding across all samples (Fig S8). Alterations in LOS composition due to these mutations could be subtle and not easily observed via this method given that rhamnose is expected to be within the LOS core, thus we sought alternative hallmarks of altered outer membrane structure. A slight sensitivity to growth in LB medium was observed for the Δ*rfbD* mutant consistent with such a defect (Fig. S9). Susceptibility to high-molecular weight antibiotics, such as vancomycin and erythromycin, has been used as a proxy for increased outer membrane permeability (39). The *A. tumefaciens rfb* mutants display increased sensitivity to vancomycin and erythromycin, with the Δ*rfbD* mutant displaying the most dramatic increases in sensitivity (Fig 8A, 8C). Increased sensitivity to high molecular weight antibiotics manifests to varying extents for the *rfb* pathway mutants, in parallel with the graded motility deficiency in each mutant. Mutants in *rfbD* appear to have the most drastically increased sensitivity, followed by Δ*rfbC* and Δ*rfbCBDA,* with only subtle differences in Δ*rfbA* and Δ*rfbB.* While individual *rfb* mutants display increased sensitivity compared to wild-type C58, suppressors isolated for their effect on motility in the nonmotile Δ*rfbD* show varying levels of suppression of the increased antibiotic sensitivity of C58 Δ*rfbD*. For example, the Δ*rfbA*Δ*rfbD* mutant displays sensitivity similar to wild type while other motility suppressor mutants display variable sensitivities, though all decrease antibiotic sensitivity to some extent in the C58 Δ*rfbD* mutant. Unsurprisingly, the ΔATU_RS21610Δ*rfbD* suppressor mutant in which motility is least rescued likewise shows the most modest rescue of the increased antibiotic sensitivity (Fig. 8B). The role of LPS in successful agrobacterial plant infection via interactions with plant hosts has been previously established (40). To test whether the *rfbCBDA* mutants are affected for virulence due to altered surface polysaccharide structure or surface attachment, plant infection assays were performed using manual inoculation of wounded *Kalanchoe daigremontiana* leaves. All *rfb* mutant strains remain pathogenic with no discernable deficiency in this qualitative assay (Fig. S10).

**Figure 8.**
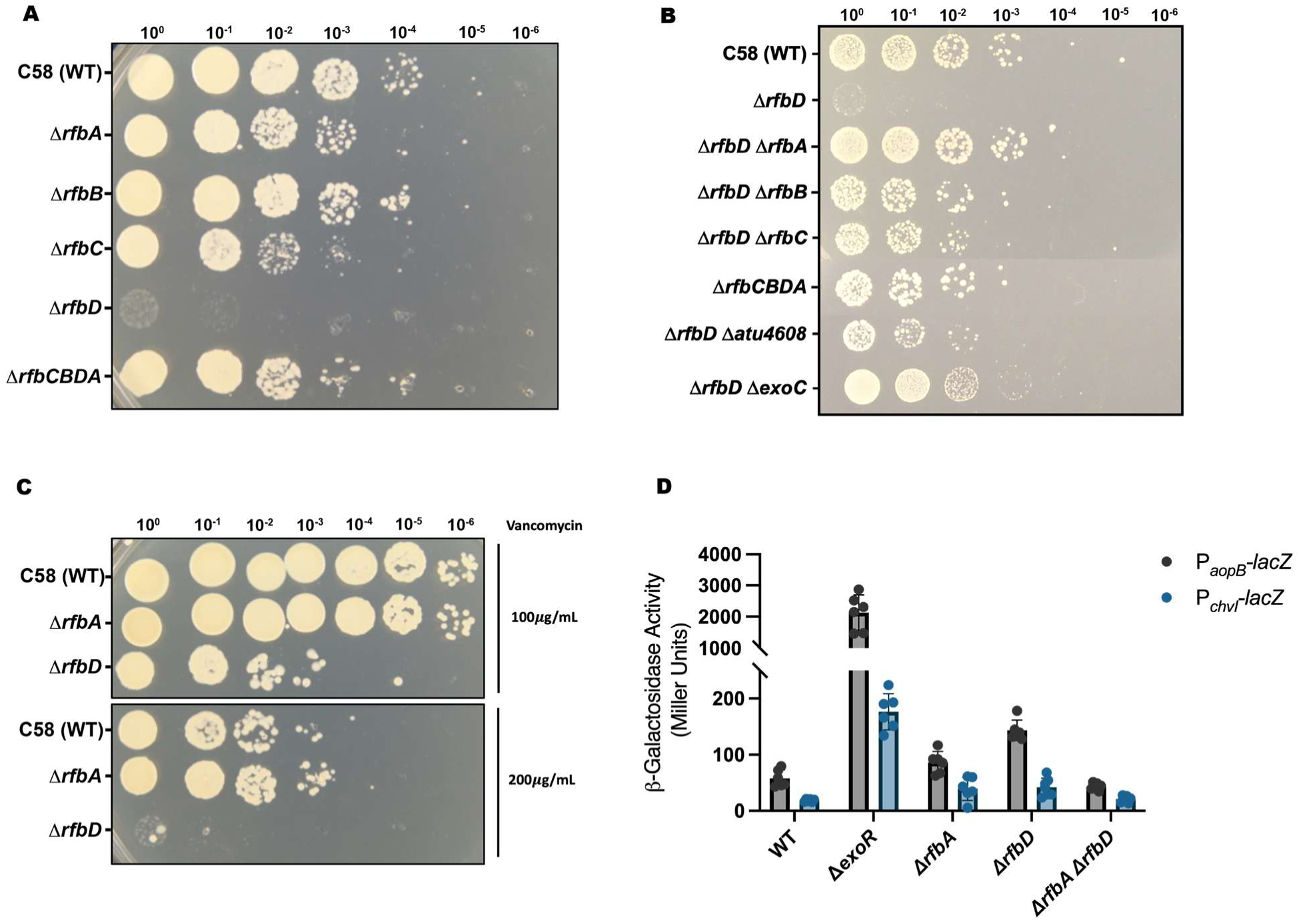
Increased susceptibility to high molecular weight antibiotics and activation of outer membrane stress signaling. (A) Colony spot dilutions of each *rfb* deletion mutant in otherwise WT C58 challenged with 100 *µ*g/mL of erythromycin in LB for 18 h prior to plating for recovery and imaging after 48 h at 30°C. B) Erythromycin (100 *µ*g/mL) challenge for motility suppressor mutant strains, performed as in A. C) Colony spot dilutions for *rfb* mutants with treatment of 100 *µ*g/mL or 200 *µ*g/mL vancomycin in LB for 18 h prior to plating for recovery and imaging after 48 h of growth at 30°C. (D) Beta-galactosidase assays of exponential phase cultures grown in ATGN broth of indicated mutants in WT C58; harboring *lacZ* fusions to ChvGI target promoters *P_aop_*_B_ and *P_chvI_*.

Outer membrane stress signaling within the *Rhizobiales* is mediated in part by the ChvG-ChvI two-component system (TCS), which is broadly conserved within the APB group (41). In *A. tumefaciens* ChvG-ChvI signaling involves the inhibitory periplasmic protein ExoR that associates with the periplasmic domain of ChvG. Under stress such as acidic conditions, ExoR is proteolytically cleaved thereby releasing ChvG to actively phosphorylate ChvI and regulate target genes (42, 43). Activation of this system in *A. tumefaciens* results in a wide range of behaviors, such as down-regulation of motility, inhibition of biofilm formation, and activation of the Type VI secretion system (T6SS) (43–45). Transcriptionally activated targets of ChvI in *A. tumefaciens* include *aopB, chvI,* and the T6SS *hcp* operon (42). We hypothesized that the increased outer membrane permeability reflected by increased antibiotic susceptibility in the *rfb* mutants, might stimulate the ChvG-ChvI pathway. To test for ChvG-ChvI activation, plasmid-borne *lacZ* transcriptional fusions for the *aopB* and *chvI* genes were introduced into *rfb* mutant backgrounds. Both the *aopB-lacZ* and *chvI-lacZ* fusions are significantly increased for β-galactosidase activity in the Δ*rfbA* and Δ*rfbD* mutants relative to wild type, though less-so relative to the Δ*exoR* mutant, in which ChvG is constitutively active (Fig. 8D). The Δ*rfbA* Δ*rfbD* double mutant that suppresses motility and antibiotic sensitivity of the Δ*rfbD* mutant also expresses these fusions at wild type levels. These observations are consistent with the perturbation of the dTDP-β-L-rhamnose pathway causing an outer membrane stress response within the cell.

## Discussion

Prior work aiming to elucidate genetic components responsible for biosynthesis of each of the two UPP species, UPP_GlcN_ and UPP_GalN_, presumptively identified several mutated genes that affected one of the two polysaccharides (14). In-depth examination of *rfbA* and *rfbD*, two genes identified specifically in the Δ*uppW* mutant disabled for synthesis of UPP_GlcN_ but still capable of UPP_GalN_ production, has validated their initial identification but also revealed unexpected complexity (14). Mutation of these genes decreases UPP production and UPP-dependent biofilm formation and causes a general increase in cellular aggregation that also requires the UPP. The increased aggregation is consistent with studies of *C. crescentus* holdfast-mediated adhesion, where mutants in LPS biosynthesis genes increased non-specific aggregation (29). Mutants in the *rfbCBDA* genes are affected for flagellar motility, and manifest outer membrane permeability defects as determined by antibiotic sensitivity, as well as activating the ChvGI outer membrane stress response.

Despite the identification of these dTDP-β-L-rhamnose precursor genes in a screen for contributors specific to the UPP_GalN_ polysaccharide species, we have shown there to be both UPP_GalN_ dependent effects on multicellular behaviors, such as attachment and auto-aggregation, as well as UPP independent changes to motility and outer-membrane stress responses. This work has further interrogated the composition of the UPP adhesin and its specific determinants, as well as the overlapping requirement of these genes in motility and outer membrane function. Our findings demonstrate how disruption of polysaccharide precursor pools can broadly impact cellular phenotypes extending to separate systems, such as that of UPP adhesin biosynthesis and outer membrane biogenesis (Fig. 9).

**Figure 9.**
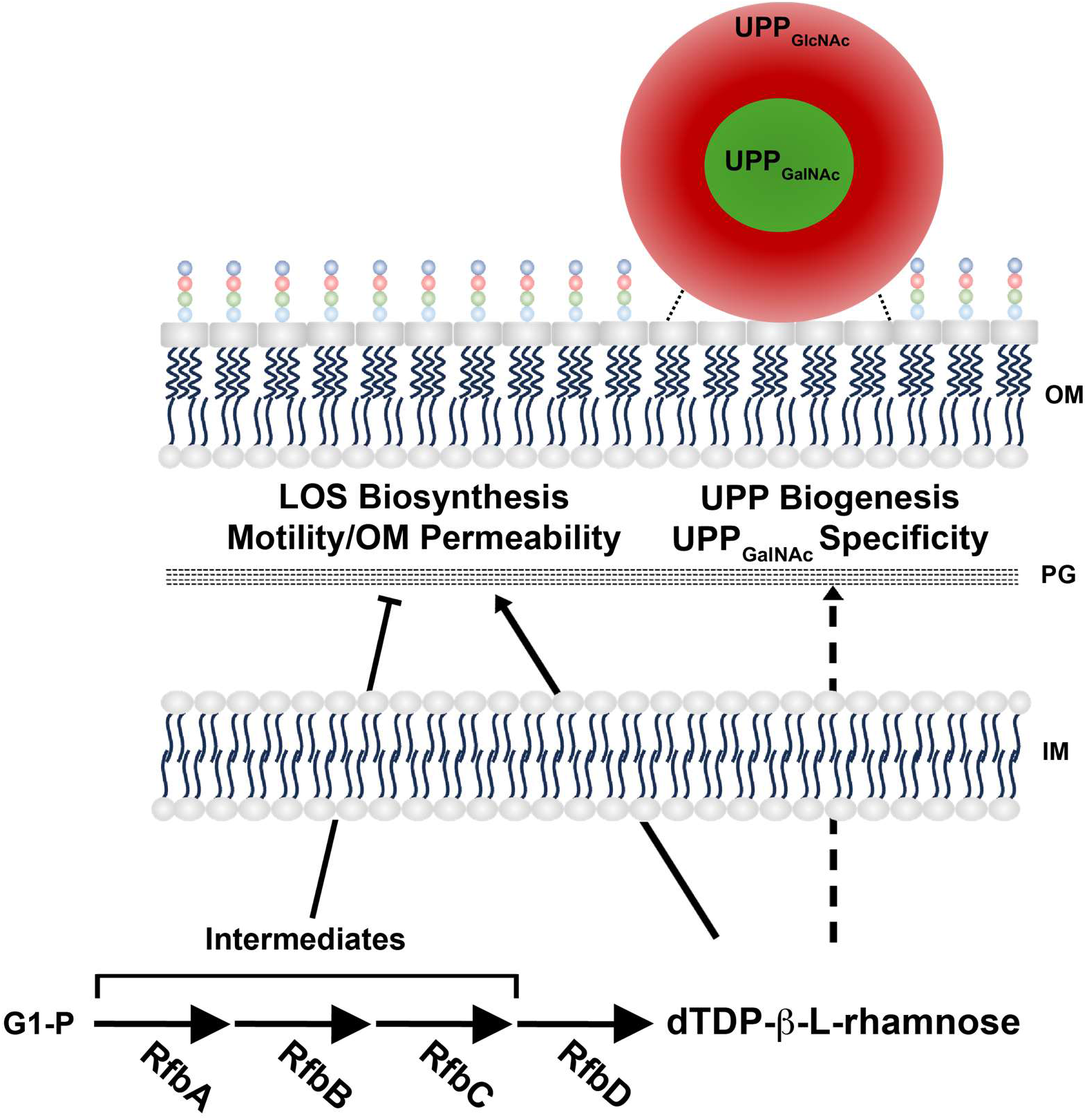
Prospective model for impacts of dTDP-B-L-rhamnose pathway on flagellar motility and UPP-mediated surface attachment. Model depicts the usage of rhamnose derivatives across cell surface functions such as UPP biosynthesis and LOS biosynthesis. The *rfb* operon encodes the biosynthetic proteins necessary for conversion of glucose-1-phosphate and deoxythymidine triphosphate (dTTP) into dTDP-β-L-rhamnose. This sugar nucleotide precursor is likely directly used for the rhamnose moieties within the core oligosaccharide of the LOS and impacts UPP biosynthesis and/or is important for its function, as well as motility and outer membrane permeability.

### Identification of *rfbA* and *rfbD* genes and their connection to UPP-dependent biofilm formation and aggregation

The *rfbCBDA* operon structure in *A. tumefaciens* suggested that the initial transposon mutants isolated in *rfbD* might have been polar on *rfbA*, raising the possibility that the DCR phenotype could have been solely due to the RfbA deficiency that normally drives the first committed step of the dTDP-β-L-rhamnose pathway. However, in-frame deletion of *rfbD* has more severe phenotypes than the *rfbA* deletion mutant, and is well complemented with plasmid-borne, ectopically expressed *rfbD*. Thus, mutation of the first step of the pathway and the final step producing dTDP-β-L-rhamnose both impact the pathway, but to different extents. In fact, the intermediate pathway enzymes encoded by *rfbB and rfbC* also influence several of the same phenotypes, including CR staining, motility and increased aggregation, but also to different levels of severity. The DCR phenotypes of these nonpolar deletion mutants verify their isolation from the UPP_GalN_ (Δ*uppW*) specific transposon screen and the biofilm phenotypes largely corroborate this data. There appear to be distinct phenotypic differences dictated by the Wzy-type polymerase present, suggesting that the presence of these genes is necessary for normal UPP production, with absence of this precursor more acutely affecting UPP_GalN_.

Mutation of *rfb* genes or their homologs in several other bacteria has been shown to decrease EPS and biofilm formation, usually due to alterations in outer membrane structure or other undefined mechanisms (24, 33, 46, 47). Availability of polysaccharide precursor pools can influence multiple different polysaccharide synthesis pathways (48). The composition of the UPP itself is under investigation, and it is plausible that dTDP-β-L-rhamnose could feed directly into this pathway. A rhamnose-specific lectin (from chum salmon, *Onchorhyncus keta*) does not label the UPP, providing preliminary evidence against the direct incorporation of rhamnose (data not shown). However, blockage of the RfbABCD pathway may influence that of a different precursor polysaccharide subunit, given that single mutants in the *rfb* genes in otherwise wild type backgrounds show increased polysaccharide production by CR staining while the corresponding mutants in the CDGS- background do not. Similarly, it is possible that loss of *rfb* genes downstream of the RfbA driven reaction leads to aberrant accumulation of polysaccharide nucleotide precursors, such as TDP-glucose, and this interferes with normal UPP synthesis. Likewise, loss of *rfbA* would decrease demand for glucose-1-phosphate, thereby increasing its levels. In either case, these intermediate metabolites could then go on to alter substrate pools for the UPP. It has been reported that dTDP-β-L-rhamnose can allosterically regulate RfbA activity via negative feedback inhibition (34, 35). If this is the case in *A. tumefaciens*, mutation of *rfbD* would abolish allosteric control of the pathway by dTDP-β-L-rhamnose and could increase entry of glucose-1-phosphate into this pathway and correspondingly alter precursor pools (49, 50).

In several cases, the decreased surface-attached biofilm formation and increased aggregation we observe are more pronounced in the Δ*uppW* and native UPP backgrounds, consistent with dependence on the UPP_GalN_ polysaccharide. Moreover, the phenotypes are less dramatic or fail to manifest in the Δ*uppY* mutants in which UPP_GalN_ is absent. These trends are in line with the initial mutant isolation in the UPP_GalN_-specific screen. In many cases, Wzy polysaccharide polymerases such as the UppY and UppW proteins can display promiscuity with different substrates (51). Changes in polysaccharide precursor pools could therefore impact downstream products by increasing interaction with non-cognate substrates, that would otherwise be too low affinity. It is unknown whether proteins of the UPP pathway display such promiscuity (14).

The *rfbCBDA* mutants manifest biofilm defects to different severities, with the loss of *rfbA, rfbB, rfbD and rfbCBDA* more pronounced in wild type C58 with relatively little impact in the CDGS- Δ*pruA* strain with high UPP production. The variable motility defects observed in these mutants likely impact surface attachment and biofilm formation, and these processes are clearly interlinked. Generally, loss of *rfbD* appears to impart much more dramatic phenotypes, with this being the only biofilm deficient mutant in the elevated UPP production (CDGS- Δ*pruA*) background. Our findings reveal some limited potential for partial rescue between paralogous genes such as *rfbA* and *exoN* (ATU_RS18920), but this does not fully explain the range of phenotypes observed. For *rfbB*, *rfbC* and *rfbD* mutants, decreases in dTDP-β-L-rhamnose would alleviate feedback inhibition of RfbA, and may result in greater glucose-1-phosphate flux into the pathway and buildup of pathway intermediates. Additionally, the anchoring of UPP to the outer membrane is not well understood. The O-antigen ligase homolog UppF may potentially function in this anchoring and impacts to LOS structure due to mutation of dTDP-β-L-rhamnose could in turn modulate anchoring of the UPP to the outer membrane (52). The aggregation and specific decreases dictated by UPP_GalN_ show differential impacts for each of the two UPP-subspecies, with exacerbated increased aggregation in strains with UPP_GalN_, which is likely reflective of the decreases in attachment, biofilm formation and CR staining in the Δ*uppW* background in which *rfbA* and *rfbD* Tn mutants were initially identified. Considering the disruption of aggregates with UppH treatment and their absence within the Wzy-polymerase double mutant, it is likely that this attachment is most strictly mediated by the UPP, rather than by changes to the outer membrane of these *rfb* mutants. In fact, the attachment and auto-aggregatory phenotypes characteristic of these mutants are likely impacted not only by changes to the adhesin but also the defects in motility and outer membrane integrity.

### Inhibition of flagellar-mediated motility

A major class of related phenotypes caused by *rfbCBDA* mutations are motility and outer membrane defects. These two phenotypes follow similar degrees of severity in specific *rfb* mutants, with the most severe effects in Δ*rfbD*, and are similarly suppressed by secondary mutations. Disruption of *rfb* genes or homologs in several other systems can result in motility defects, perhaps through a compromised outer membrane that interferes with normal flagellar rotation (28, 33, 34). Our findings reveal substantial motility defects across a range of severity for individual *rfbCBDA* mutants in *A. tumefaciens*. The Δ*rfbD* mutant is largely nonmotile, with Δ*rfbC* next most dramatically affected, followed by Δ*rfbB* and Δ*rfbA*. The *rfbCBDA* gene mutants, including the non-motile Δ*rfbD* mutant, produce visible flagella with impaired function (Fig 5B and data not shown). Thus, the motility defect appears to be due to decreased or abolished flagellar rotation. Conceivably, the elevated production of UPP that results in greater aggregation might impede flagellar rotation, but incubation of motility assays in the presence purified UppH which cleaves both UPP species and decreases surface attachment, does not improve mutant motility (data not shown). Furthermore, deletion of *uppY* and *uppW* in the CDGS- Δ*pruA*Δ*rfbD* non-motile mutant in which UPP production is abolished, does not rescue motility. However, in the wild type strain background the Δ*uppW* mutation suppresses the Δ*rfbD* motility defect, whereas the *uppY* mutation does not. The basis for this Δ*uppW* suppression remains unclear, but it is striking that in a non-biased Δ*rfbD* suppressor screen in the otherwise wild type C58 background, no mutations were obtained in *uppW* despite the identification of multiple mutants in the *rfbA, rfbB*, *rfbC* and *exoC* genes.

Motility in *A. tumefaciens* is controlled by the master regulators VisN and VisR, which activate expression of the response regulator *rem*, the product of which in turn activates expression of the flagellar biogenesis genes (15). The ChvG-ChvI system activates expression the MirA protein which is an anti-activator of Rem and can block motility (42). Therefore, given that mutation of *rfbA and rfbD* stimulate ChvG-ChvI target gene expression, activation of *mirA* expression could inhibit motility. However, the non-motile *rfbD* mutants still make flagella, suggesting that there is no block to assembly, as would be expected for elevated MirA. Furthermore, ectopic expression of Rem from *P_lac_-rem* constructs in the Δ*rfbD* mutant fails to improve motility (data not shown), suggesting that ChvG-ChvI dependent inhibition of Rem through *mirA* is not responsible for the non-motile phenotype. Overall, the simplest model to explain the motility phenotypes of the *rfbC*, *rfbB, rfbA*, and *rfbD* mutants is that these mutations differentially affect the level of dTDP-β-L-rhamnose (Δ*rfbD>*Δ*rfbC*>Δ*rfbB*>Δ*rfbA*) and its precursors, perhaps through partial rescue by pathway paralogs as with *rfbA* and *exoN* (ATU_RS18920), and the decrease in this LOS precursor results in outer membrane perturbation that impedes flagellar rotation.

However, the simple model does not readily explain why the Δ*rfbD* mutant is so profoundly blocked for motility. Furthermore, it is striking that this mutant is largely, although not completely suppressed for its severe motility deficiency by mutations that disrupt earlier steps in the pathway. Similarly, deletion of the phosphoglucomutase gene *exoC* that provides glucose-1-phosphate precursor to the RfbA enzyme, partially suppresses the Δ*rfbD* mutation for motility. These observations suggest that in the Δ*rfbD* mutant, the dysregulation of precursor metabolites into this pathway lead to even greater severity in its motility defects. As mentioned above, the absence of dTDP-β-L-rhamnose would lead to loss of feedback inhibition, and greater RfbA-mediated entry of G-1-P and dTDP into the pathway. Accumulation of an inhibitory pathway intermediate would therefore be maximally elevated. As the Δ*rfbC* mutation causes effective suppression of Δ*rfbD*, this implicates its product of 4-keto-dTDP-β-L-rhamnose as a strong candidate for an inhibitor. It is conceivable that at elevated levels in the absence of dTDP-β-L-rhamnose, 4-keto-dTDP-β-L-rhamnose could be incorporated into LOS, or might inhibit its normal assembly, and create even more profound outer membrane defects. This model is also supported by the observation that the entire operon deletion, including *rfbD*, has the equivalent motility phenotype as Δ*rfbA*. It would manifest the lack of dTDP-β-L-rhamnose, but not the exacerbation of the RfbA, RfbB and RfbC-generated intermediate. Ectopic expression of *rfbCBA* in the operon mutant results in decreased motility, further demonstrating that the accumulation of pathway intermediates accentuates the already deficient motility resulting from a loss of dTDP-β-L-rhamnose. The suppression of the Δ*rfbD* motility defect with transposon insertions in *exoC* is also interesting given that nonpolar deletion of *exoC* itself leads to motility defects, decreased biofilm formation, and decreased Congo Red staining (14). The *exoC* single mutant would be similar to mutations in the RfbABCD pathway leading to a decrease of flux through this pathway, thereby rescuing the defect of the Δ*rfbD* mutant.

The only gene identified not known to be involved in the dTDP-β-L-rhamnose pathway is that of ATU_RS21610, a dual domain protein with methyltransferase and glycosyltransferase domains. Using AlphaFold predicted models, the methyltransferase domain of this protein displays homology to crystal structures of methyltransferases involved in NDP-N-methyl-L-glucosamine biosynthesis (*B. cereus)*, while the glycosyltransferase domain displays homology to RmlT (*Listeria monocytes*) and WTA β-O-GlcNAcylation (*Staphylococcus aureus*) (Foldseek PDB search (53)). The homology of ATU_RS21610 to other proteins that function in polysaccharide precursor biosynthesis is similar to the other Δ*rfbD* suppressors, consistent with its requirement for the Δ*rfbD* imposed motility defect. However, mutation in ATU_RS21610 alone in the otherwise wild type has no effect on motility.

### Outer membrane deficiencies and stress signaling

Mutations that compromise outer membrane structure are well known to increase the permeability of this barrier to large molecules (39, 54, 55). The observed increased sensitivity to large molecular weight antibiotics in the *rfbCBDA* mutants is consistent with outer membrane permeability defects. Rhamnose is often incorporated into the LPS/LOS of many bacteria, predominantly in the O-antigen (56, 57), although studies in *A. tumefaciens* suggest it is incorporated into the core component (22, 58). Loss of rhamnose incorporation into the core or O-antigen could lead to permeability defects in the outer membrane, and this has been observed in other bacterial systems in which the dTDP-β-L-rhamnose pathway has been mutated (24, 27, 59). The degree of antibiotic sensitivity in the *rfbCBDA* mutants follows the same trend as the motility defects, with Δ*rfbD* the most sensitized, followed in order by Δ*rfbC*, with Δ*rfbA* and Δ*rfbB* exhibiting far less sensitivity. The suppressor mutations isolated based on motility restoration in the Δ*rfbD* mutant diminished the increased antibiotic sensitivity, and they also improve UPP-dependent biofilm formation in this mutant. The correlation of motility deficiencies, increased antibiotic sensitivities, and biofilm formation, as well as subsequent suppression of each phenotype in transposon mutants isolated for restored motility strongly suggests strongly that these phenotypes are connected. This is most likely through changes in outer membrane composition imparted by mutations to the *rfb* operon. It could be that the buildup and/or shunting of intermediate *rfbCBDA* metabolites to other metabolic networks could be responsible for this perturbation, as the Δ*rfbA*Δ*rfbD* double mutant both suppresses motility and outer membrane defects as well as prevents activation of ChvGI. Furthermore, activation of the ChvG-ChvI TCS implicates perturbation of the outer membrane composition that is activating stress signaling (60). Indeed, mutations in LPS precursor genes has been shown to activate the outer membrane stress signaling Rcs pathway in *Enterobacteriaceae*, also leading to defective motility (61, 62) The increases in activity of ChvG-ChvI target genes in *A. tumefaciens* suggests activation of this system, and again are indicative of deficiencies in the outer membrane. However, ChvG-ChvI activation blocks flagellar gene expression and assembly, and Δ*rfbD* does make flagella but cannot use them to swim. Therefore, this regulatory pathway may contribute to the complex phenotypes observed in the *rfbD* mutant but is most likely not the cause of the motility defect.

In summary, this work demonstrates the multifaceted impact that the *rfbCBDA* operon has in supplying the dTDP-β-L-rhamnose precursor for the outer membrane and other bacterial cell surface structures. Perturbation of this pathway can have effects that exceed simply the loss of this precursor, and the greater susceptibility to antibiotics manifested by pathway mutants suggest that targeted approaches could augment antimicrobial therapies and modulate complex bacterial activities such as motility and biofilm formation.

## Materials and Methods

### Strains, reagents, and bacteriological methods

Bacterial strains, plasmids, and the oligonucleotides used in this study are listed in supplementary tables S2 and S3, respectively. *A. tumefaciens* strains were grown at 25°C or 30°C on liquid or solid LB (10 g Bacto-tryptone, 5 g yeast extract, 5 g NaCl per liter, pH 7.0) or AT minimal medium (63) with 0.5% (w/v) glucose and 15 mM ammonium sulfate (ATGN). When performing biofilm assays, 22 µM FeSO_4_ was added into the liquid ATGN media immediately prior to inoculation, and for *sacB* counter-selection, 2.5-5% (w/v) sucrose replaced glucose as the sole carbon source to make ATSN. For ATGN-CR, the CR dye was dissolved in Milli-Q water (Millipore Sigma, Burlington, MA) to a concentration of 20 mg/ml, then passed through a 0.2 µm filter to remove aggregates. Congo Red phenotypes were assayed on 75 ug/mL ATGN-CR. *E. coli* strains were grown at 37°C in LB medium supplemented with appropriate antibiotics.

Plasmids were introduced into chemically competent cell preparations of *E. coli*, and into *A. tumefaciens* via conjugation or electroporation (64). Oligonucleotide primers were obtained from Integrated DNA Technologies, Coralville, IA, and Sanger sequencing was performed by ACGT, Inc., Wheeling, IL. Chemicals, antibiotics, and culture media were obtained from Fisher Scientific and Sigma-Aldrich. When necessary, antibiotics were added to the medium as follows: 100 µg/ml ampicillin (Ap), 50 µg/ml gentamycin (Gm) and 50 µg/ml kanamycin (Km) for *E. coli*, then 500 µg/ml Streptomycin, and 300 µg/ml Gm and Km for *A. tumefaciens*. Isopropyl-β-D-thiogalactopyranoside (IPTG) was added to 500 μM to induce expression of plasmid borne genes.

### Plasmid construction

DNA sequences were amplified from *A. tumefaciens* C58 genomic DNA and cloned into corresponding vectors using a combination of Gibson Assembly (65) and Fastcloning approaches (66). Specifics for plasmid construction are provided in the Supplemental Methods.

### Construction of in-frame, marker-less deletions

Generally, In-frame marker less deletions were constructed as previously described (19, 64) with full specific details provided in the Supplemental Methods. Upstream and downstream ∼ 500 bp regions flanking the genes to be deleted were amplified from *A. tumefaciens* C58 genomic DNA using specific primers (P1 and P2 for upstream regions, P3 and P4 for downstream regions) and Q5 polymerase. Care was taken to ensure the 5’ and 3’ ends of the genes were not altered, and that translational coupling between linked genes was still retained. When necessary, primers P2 and P3 were designed with reverse complementarity in the 5’ sequences to allow for single overlap extension (SOE) amplification of both genes into a single amplified product as previously described (64). The combined flanking regions were fused with the appropriately linearized suicide vector pNPTS138 using Fast cloning or Gibson assembly to create a deletion construct which was subsequently transformed into chemically competent S17-1/λpir *E. coli*. The pNPTS138 vector confers kanamycin resistance (Km^R^) and sucrose sensitivity (Suc^S^) due to the presence of the *sacB* gene. Introduction of this plasmid into *A. tumefaciens* was via conjugation with S17-1/λpir *E. coli*. The ColE1 origin of pNPTS138 does not replicate in *A. tumefaciens*, and a single crossover event allows plasmid integration into the chromosome which is then confirmed by patching transformants on to ATGN-Km and ATSN to screen for Km^R^ Suc^S^ strains. Strains that have excised the plasmid were then isolated by parallel patching on to ATSN and ATGN-Km to screen for Km^S^ Suc^R^ derivatives. Deletion of the target genes is confirmed by diagnostic PCR using primers (P1 and P4) flanking the site of deletion. Complementation strains were generated by electroporating the indicated pSRKGm derivative (67) into the corresponding deletion mutant, while vector control strains were generated likewise with the empty pSRKGm plasmid.

### Motility Assays

Motility assays were performed in 0.25% ATGN agar. When enriching for motility suppressors, 5 μL aliquots of the transposon library were inoculated in the center of the motility agar plate and grown at room temperature for 48-72 hr. When testing motility signatures otherwise, fresh streaks of strains grown on ATGN or LB were used to inoculate the center of motility agar plates by tapping the surface of a colony with inoculating needle or toothpick and inoculating directly into the center of the plate.

### Congo Red assays

*A. tumefaciens* strains were grown in ATGN until mid-exponential phase, normalized to an OD_600_ of 0.05, and 3 µl spotted on to ATGN-CR plates with or without 500 μM IPTG. Plates were incubated at 30°C. Photographs of spots were taken following incubation for 48 h.

### Biofilm assays

Strains of *A. tumefaciens* were grown and analyzed as previously described (64). PVC coverslips were inserted vertically into the wells of 12-well polystyrene plates (Corning Inc.) and UV sterilized for 20 minutes. The wells were filled with 3 ml mid-exponential phase cultures (with or without IPTG) normalized to an OD_600_ 0.05. The 12-well plates were incubated in static conditions for 48 h at room temperature inside a plastic Tupperware container with an open bottle of saturated K_2_SO_4_ solution for maintaining humidity. Following incubation, the coverslips were rinsed with sterile water and stained with 0.1% w/v crystal violet (CV) solution 0.1% w/v for 10 min. Stained coverslips were rinsed with sterile water and then submerged in weigh boats containing 1 ml 30% acetic acid to solubilize the CV. Quantification was performed by measuring soluble A_600_ CV absorbance using a Biotek Synergy HT microplate reader and Gen 5 software (Biotek, Winooski, VT). Values were normalized to culture growth by dividing by final culture OD_600_ when biofilm scores are reported, otherwise CV A_600_ itself is reported as a measure of attachment in strains that aggregate heavily to avoid confounding effects from the lower OD_600_ in these strains.

### LOS extraction

LOS extraction was performed by an established approach (68). Briefly, cultures were grown to mid-exponential phase and washed twice in PBS before resuspension in lysis buffer. Samples were boiled at 100°C for 10 minutes before cooling at RT. A 30 µL aliquot of 2.5 mg/mL (in lysis buffer) of proteinase K was added to 150 𝑢𝑢L of cells and samples were incubated at 37°C for 24 h. LOS was then precipitated out of samples by adding 1/10 volume of 3 M sodium acetate and 2 volumes of 100% ethanol and incubating at-80 °C for 1 h. Samples were then centrifuged at 15,000 x g for 5 min, washed twice with 70% ethanol, and resuspended in sterile water to 180 𝑢𝑢L. LOS samples were visualized via the Bio-Rad silver-staining kit (#1610443).

### Antibiotic susceptibility assays

A 2 mL starter culture was grown in LB overnight for each tested strain before subculturing the following day to an OD_600_ 0.05 in the presence of the indicated concentration of antibiotic or in media alone. After an exposure period of 18 h, a 5 μL spot dilution series was then plated for recovery on LB agar. Plates were then incubated at 30°C overnight and pictures taken the following day.

### Flagellar staining and visualization

Strains to be visualized were introduced with the pBM205 [pSRKKm::com (FlaAT213C +*P_flaA_* native promoter)] (35) reporter plasmid by electroporation. Strains with pBM205 were grown overnight with 500 μM IPTG to exponential phase before incubating with 50 μM maleimide dye before fluorescence visualization on the FITC channel (filter, wavelength, scope). As an alternate technique, 10 μl of a chemical flagella stain reagent (64) was added to 3 μL of cells sandwiched between a coverslip and glass slide for microscopy. Dye was administered by capillary action, and cells were incubated 15 min prior to visualization by phase contrast microscopy on the same microscope as previously described.

### Beta-galactosidase Assays

Beta-galactosidase assay protocols were adapted from (69). Strains were grown overnight in minimal media (unless otherwise specified) until exponential phase (OD_600_ 0.4-0.8). A mixture of Z-buffer, chloroform, and 0.05% SDS was added to cells before vortexing 10 sec to lyse cells. A 100 μL aliquot of 4 mg/mL ONPG was used to start β-galactosidase reactions, and upon development of visible yellow color or extended period of incubation reactions were stopped with 600 mL of Na_2_CO_3_. Samples were then centrifuged at 16,000 x g for 3 min and the A_420_ of the supernatant was then measured. Miller units are calculated according to MU = 1000 * A420 / (OD600 * t * f), where f = volume of cells/volume of cells + volume of Z-buffer.

### Plant Infection Assays

Seedlings of *Kalanchoe daigremontiana*.were grown in greenhouse for 3 weeks prior to repotting individual plants. *A. tumefaciens* strains to be tested were grown fresh on standard ATGN solid media. *K. diagremontiana* leaves were scored using a sterile wooden stick, and each strain was then inoculated into the wounded plant tissue by smearing culture with a sterile wooden stick. Multiple leaves on multiple plants were used to test a given strain. After inoculation, leaves were then covered in plastic wrap and taped shut for 72 h to increase humidity. Plants were then grown and watered daily for 2-3 weeks until positive controls showed tumor formation.

## Supporting information

Compiled Supplemental Materials

## Acknowledgements.

This work was supported by the National Science Foundation through grant MCB 2441717 to CF, and the GEMS Biology Integration Institute, funded by the NSF DBI Biology Integration Institutes Program, Award #2022049.

## Notes

### Competing Interest Statement

The authors have declared no competing interest.

### Summary of Updates

Funding Information Supplementary Materials that were not harvested from the Journal of Bacteriology automatic submission

