## Supplementary material for "Lipopolysaccharide Precursor Mutants Disrupt Unipolar Polysaccharide Adhesin Synthesis and Cell Surface Functions in *Agrobacterium tumefaciens*": Compiled Supplemental Materials

Running title: LPS biosynthetic intermediates affect biofilms and motility

1) Supplementary Figures and Legends– S1-S10

2) Supplementary Tables – S1-S3

3) Supplementary Methods

4) Supplementary References

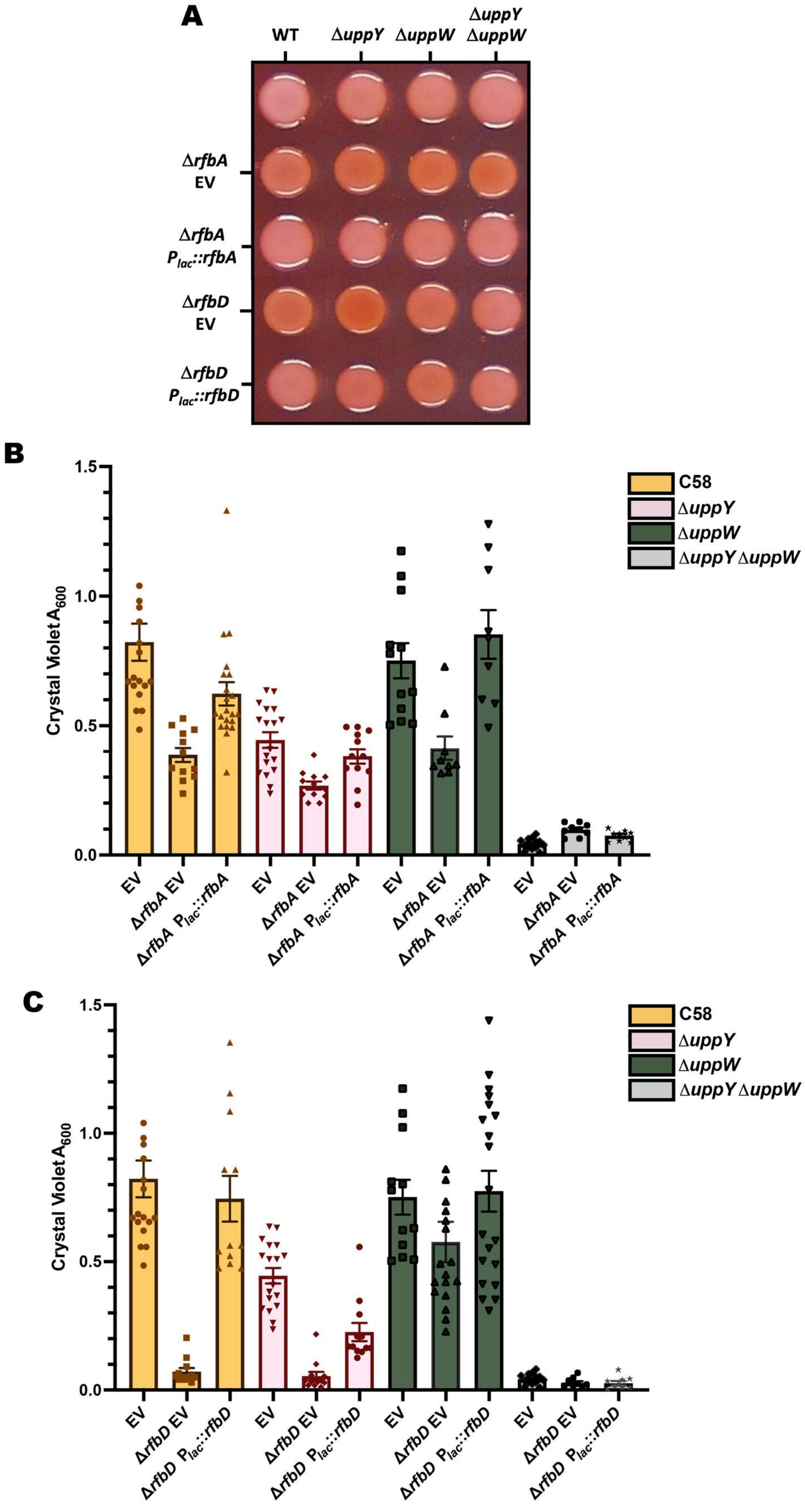

A
WT
ΔuppY
ΔuppW
ΔuppY ΔuppW
ΔrfbA EV
ΔrfbA Plac::rfbA
ΔrfbD EV
ΔrfbD Plac::rfbD
B
Crystal Violet A600
1.5
1.0
0.5
0.0
C58
ΔuppY
ΔuppW
ΔuppY ΔuppW
EV
ΔrfbA EV
ΔrfbA Plac::rfbA
C
Crystal Violet A600
ΔrfbD EV
ΔrfbD Plac::rfbD

**Figure S1: UPP-Specific and General Attachment Defects in C58 Δ*rfbA* and C58 Δ*rfbD* Mutants**. (A) CR staining of each *rfbA* and *rfbD* mutant in WT C58 with polymerase mutations. Cultures were spotted onto ATGN-CR (75 μg/mL) plates with 500 μM IPTG for 48 h. (B) CV staining of adhered cells was solubilized in 30% acetic acid and the $A_{600}$ was measured for parent strains and *rfbA* mutants in WT C58, C58 Δ*uppY*, C58 Δ*uppW*, and C58 Δ*uppY* Δ*uppW*. EV refers to the empty pSRKGm backbone while plasmid-borne copies of *rfbA* were under the control of $P_{lac}$ and induced with 500 μM IPTG. (C) $A_{600}$ CV staining for the same Wzy polymerase mutants in a WT C58 background with loss or complementation of *rfbD.* Experiment was performed as in panel B. In panels B and C, bars are the mean of biological triplicate assays, and the error bars are SEM.

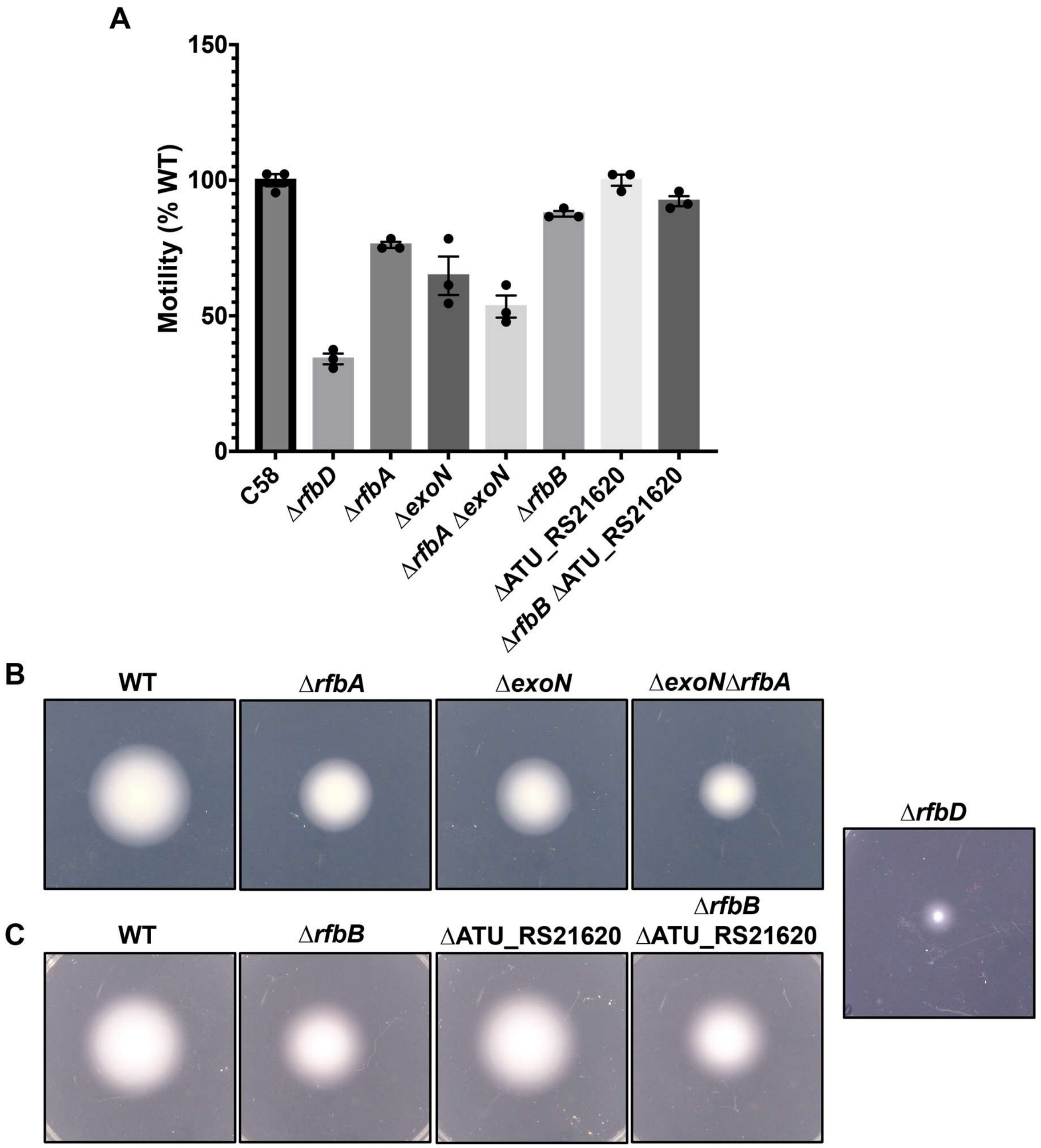

**Figure S2. Loss of *exoN* decreases motility in Δ*rfbA* mutant.** (A) Swim ring diameter measurements in WT C58 backgrounds with *rfbA*, *rfbD, exoN* and *atu4610* mutants at 72 h post inoculation of strains at RT during motility assay. Points represent mean of samples measured in triplicate with error bars representing standard deviation. (B) Representative images of mutants during motility assay. A single colony for each sample was inoculated into 0.25% agar ATGN and incubated at RT for 72 h prior to imaging.

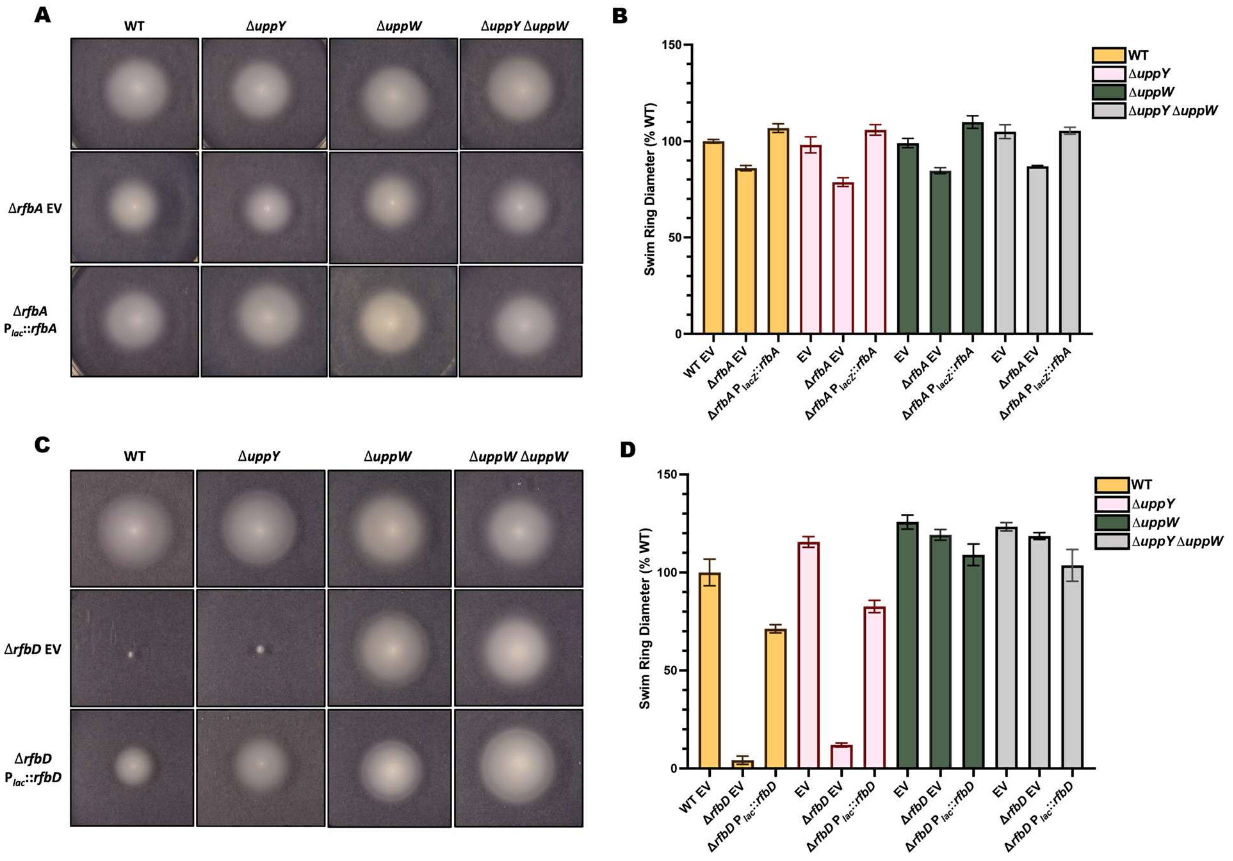

**Fig S3. Motility defects of Δ*rfbA* and Δ*rfbD* mutants in UPP polymerase mutants.** (A) Motility assays on ATGN agar (0.25%) of Δ*rfbA* mutants in WT C58 with additional *uppY* and *uppW* Wzy-type polymerase mutations. Single colonies were inoculated and incubated at RT for 3 days prior to imaging. (B) Swim ring diameter measurements of motility assays in panel A for the indicated mutants after 72 h at RT. (C) Motility assays on ATGN agar (0.25%) of Δ*rfbD* mutants with additional *uppY* and *uppW* Wzy-type polymerase mutations. Samples were inoculated and grown as in panel A. (D) Swim ring diameter measurements in C58 Wzy-mutant backgrounds with Δ*rfbD* mutants after 72 h incubation at RT. Plasmid-borne *rfbA* and *rfbD* under control of $P_{lac}$ was induced with 500 $\mu$M IPTG when indicated. EV, empty pSRKGm vector. For panels B and D, bars represent mean of samples measured in triplicate with error bars represent SEM.

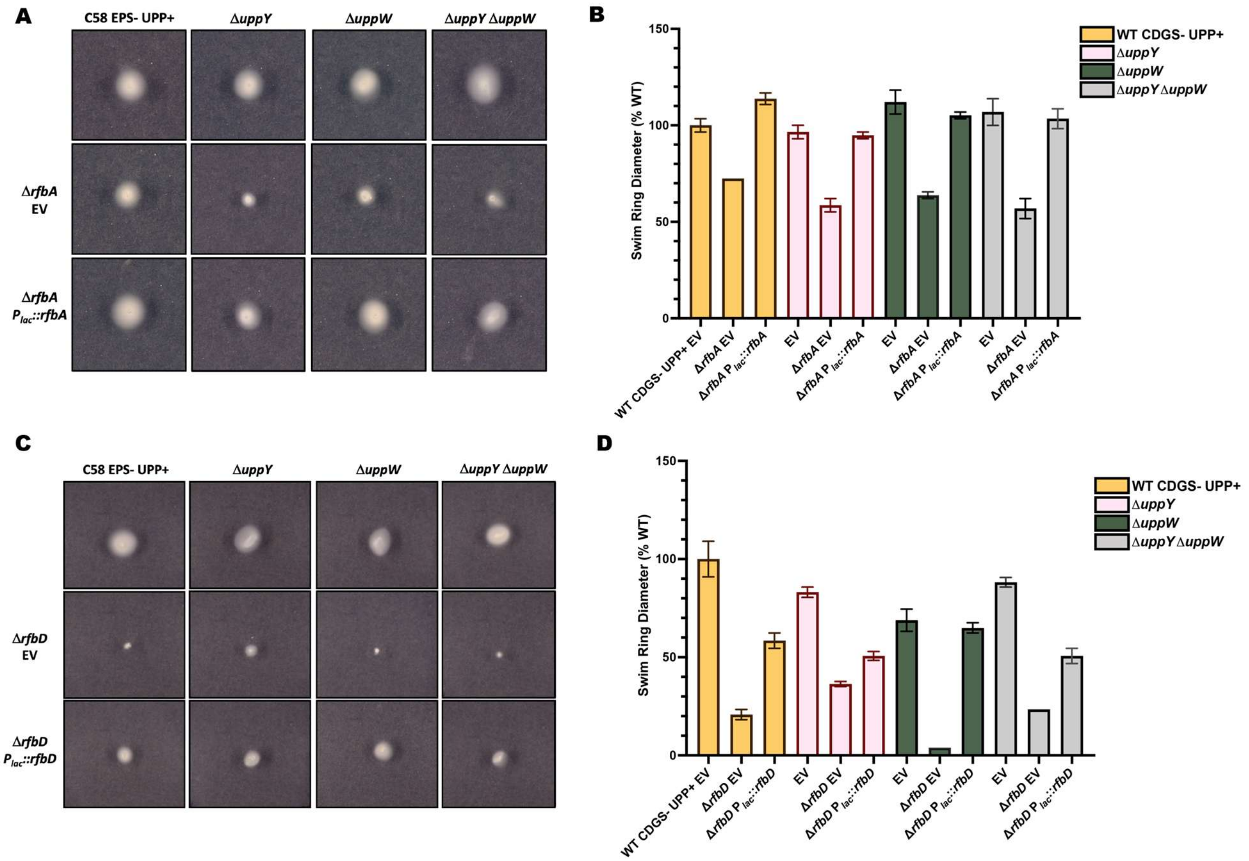

**Figure S4: Motility defects of Δ*rfbA* and Δ*rfbD* mutants in the C58 CDGS- Δ*pruA* mutant.** (A) Motility assays on ATGN agar (0.25%) of Δ*rfbA* mutants in the C58 CDGS- Δ*pruA* mutant with additional *uppY* and *uppW* Wzy-type polymerase mutations. Single colonies were inoculated and incubated at RT for 3 days prior to imaging. (B) Swim ring diameter measurements of motility assays in for the indicated mutants after 72 h at RT. (C) Motility assays on ATGN agar (0.25%) of Δ*rfbD* mutants in the C58 CDGS- Δ*pruA* mutant with additional *uppY* and *uppW* Wzy-type polymerase mutations. Samples were inoculated and grown as in panel A. (D) Swim ring diameter measurements of the mutants from panel C over 72 h incubation at RT. Plasmid-borne *rfbA* and *rfbD* under control of $P_{lac}$ was induced with 500 $\mu$M IPTG when indicated. EV, empty pSRKGm vector. For panels B and D, bars represent mean of samples measured in triplicate with error bars represent SEM.

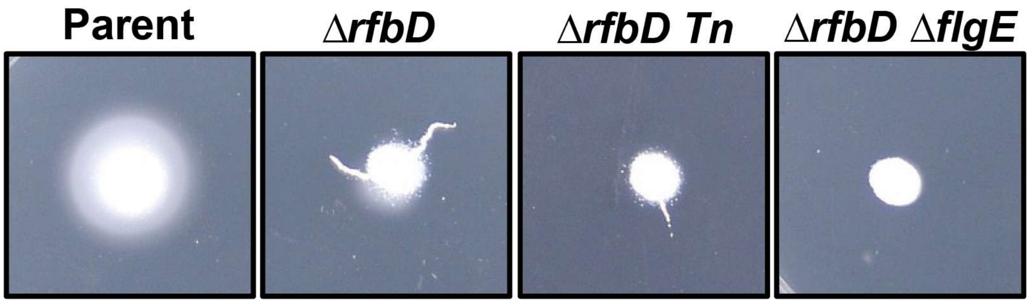

**Figure S5. Flagellar-mediated tendrils in C58 CDGS- Δ*pruA* background.** Examination of the dendritic growth pattern observed after long growth periods (72 h or longer) in the Δ*rfbD* mutant and the Δ*rfbD* transposon mutant library in the C58 CDGS-Δ*pruA* background. The tendrils that extend from the site of inoculation are on the surface of the agar and are absent with aflagellate Δ*flgE* mutants. These tendrils are distinct from the swim rings that develop due to Δ*rfbD* suppression.

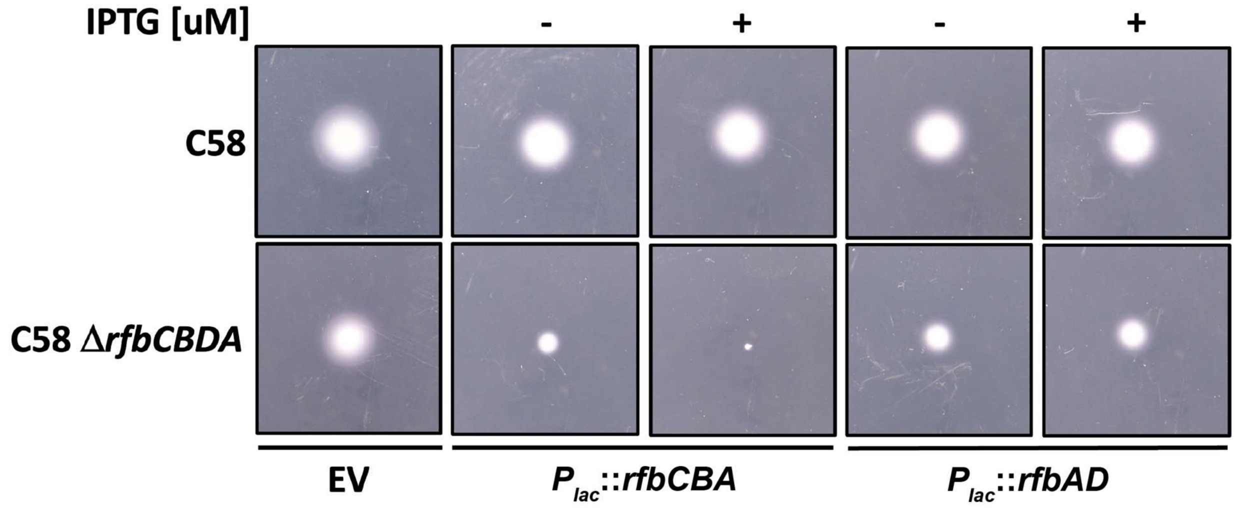

**Figure S6. Ectopic expression of early dTDP-β-L-rhamnose pathway genes can directly inhibit motility.** Motility assays on ATGN agar (0.25%) 72 h post inoculation. Plasmid-borne $P_{lac}$-*rfbCBA* and $P_{lac}$-*rfbAD* fusions are induced with with 500 μM IPTG (+) where indicated. EV, empty pSRKGm vector.

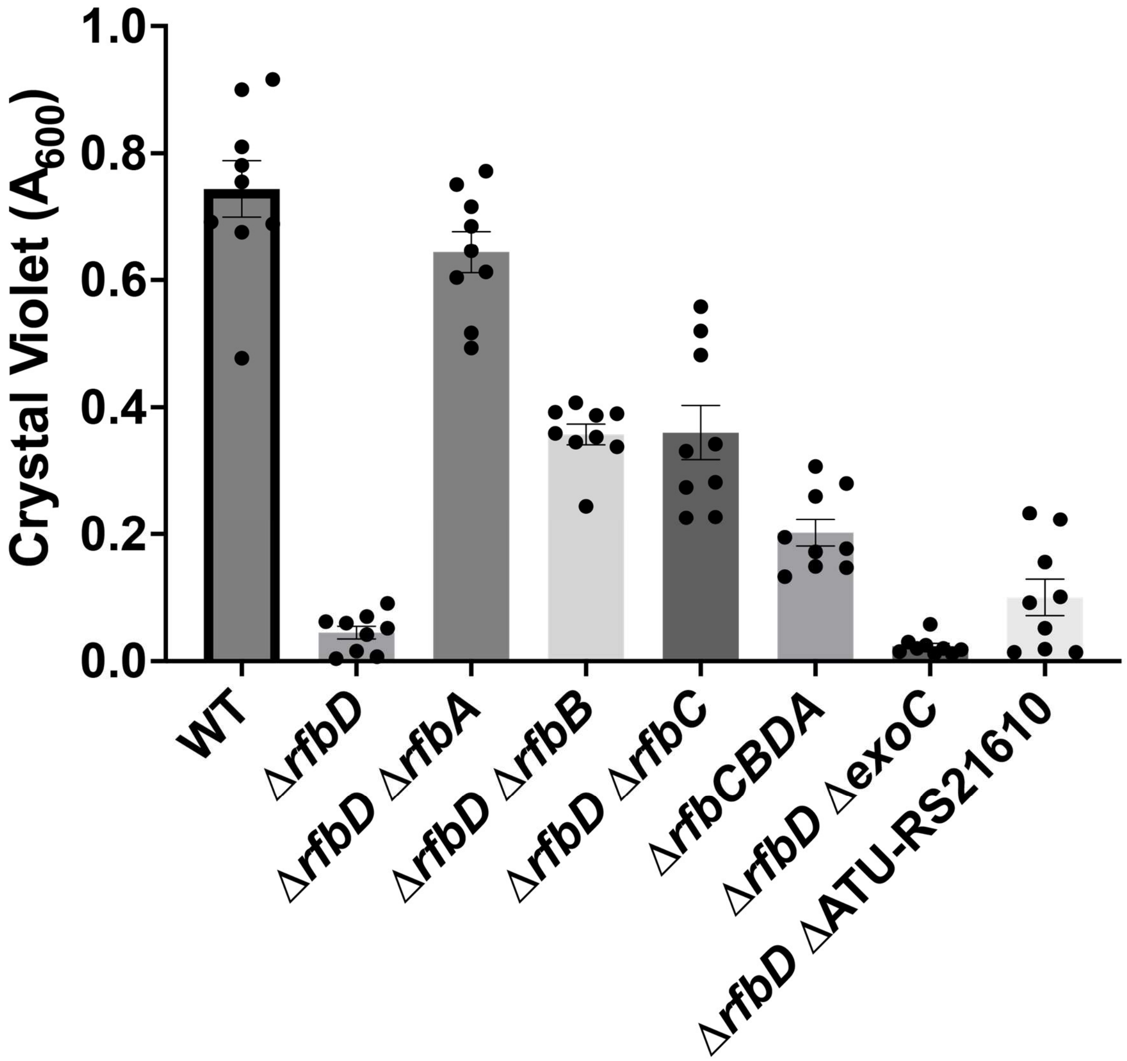

**Figure S7. Biofilm phenotype of Δ*rfbD* suppressors.** A) Biofilm phenotypes of various motility suppressors of Δ*rfbD.* CV staining of adhered cells was solubilized in 30% acetic acid and $A_{600}$ measured for each strain. Variable rescue of crystal violet attachment can be observed across strains. Bars represent the mean of biological triplicate assays and error bars are the SEM.

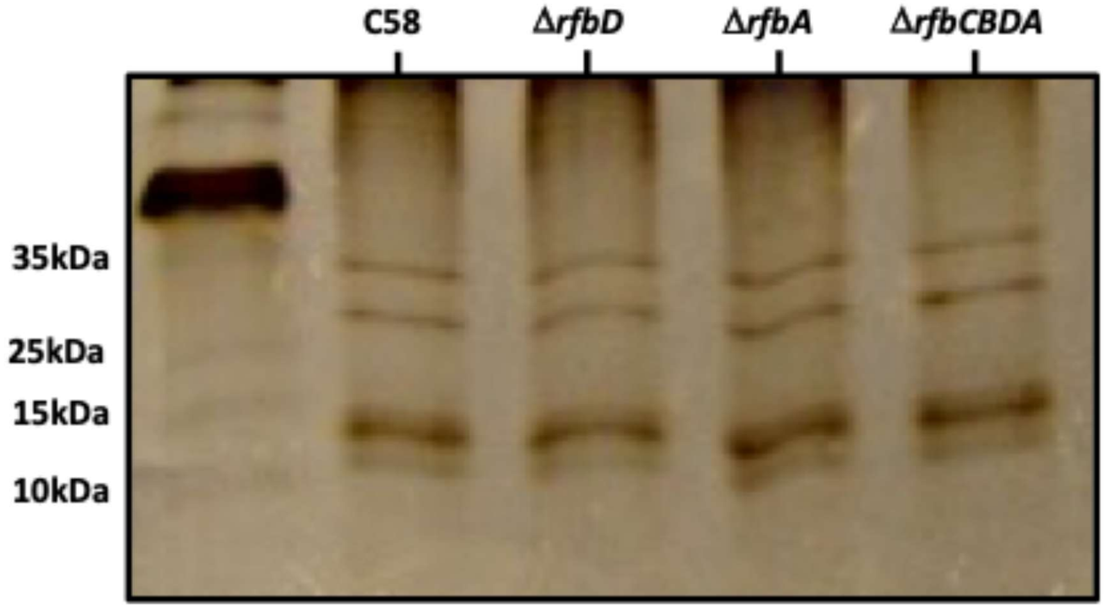

**Figure S8. LOS isolation and visualization of *rfb* mutants.** PAGE gel of LOS samples. LOS of each designated strain was extracted as described in Materials and Methods. Samples were run on a 12% 37.5:1 acrylamide:bis-acrylamide gel at 100 V for 3 h. Extracted species on the gel was then visualized using a Bio-Rad Silver-Stain kit. Lane 1 contains the NEB unstained protein ladder.

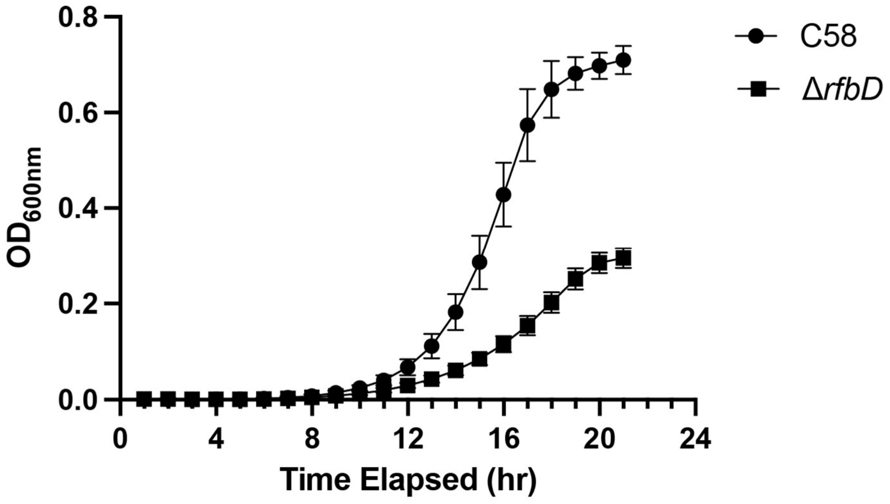

**Figure S9. Growth defect of C58 Δ*rfbD* mutant in LB broth.** Growth curve of C58 and *rfbD* mutant grown in Luria-Broth (LB). Samples were inoculated at a starting $OD_{600nm}$ of 0.01 and grown for approximately 22 h with shaking at 30°C in a Biotek Cytation plate reader.

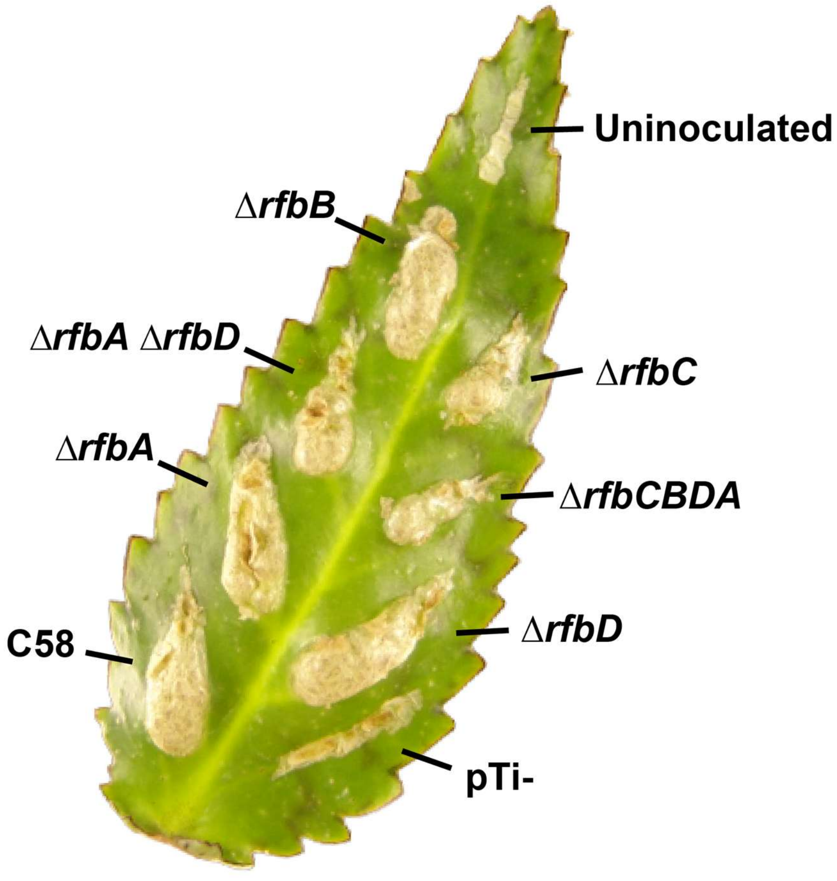

**Figure S10. *Kalanchoe daigremontiana* virulence assay of *rfb* mutants.** All otherwise WT C58 derivatives with mutations as indicated. A pTi cured mutant shows inoculation without tumor formation while the uninoculated control shows the result of only wounding. Strains were inoculated into scratch sites using growth from solid ATGN agar plates and incubated on leaves in the greenhouse ~3 weeks prior to imaging. Glass surface of image background was removed in Adobe Photoshop.

**Table S1**. Transposon insertions for motility suppression of C58 Δ*rfbD* and C58 CDGS-Δ*pruA*Δ*rfbD*

| Gene | Annotation | Insertion # C58 Δ*rfbD* Screen[1] | Insertion # C58 CDGS- Δ*pruA* Δ*rfbD* Screen[1] | Suppression in background with targeted deletion |
|---|---|---|---|---|
| *rfbA* (*atu4615*) | glucose-1-phosphate thymidylyltransferase | 3 | 8 | C58, C58 CDGS-Δ*pruA* |
| *rfbB* (*atu4617*) | dTDP-glucose 4,6-dehydratase | 5 | 2 | C58, C58 CDGS-Δ*pruA* |
| *rfbC* (*atu4618*) | dTDP-4-dehydrorhamnose 3,5-epimerase | 2 | 4 | C58, C58 CDGS-Δ*pruA* |
| *exoC* (*atu4074*) | alpha-D-glucose phosphate-specific phosphoglucomutase | 0 | 5 | C58, C58 CDGS-Δ*pruA* |
| *atu4608* | methyltransferase domain-containing protein | 0 | 2 | C58, C58 CDGS-Δ*pruA* |
| *uppW* (*atu2356*) | O-antigen ligase family protein | 0 | 0 | C58 |
| *adhP* (*atu0626*) | alcohol dehydrogenase AdhP | 2 | 0 | C58* |

[1]Independent transposon insertions isolated in this mutant background

**Table S2. Strains and plasmids used in study**

| Strain or plasmid | Relevant features | Source |
|---|---|---|
| ***E. coli*** | | |
| S17-1/λ*pir* | λ-pir, Tra+ donor for plasmid conjugation, $Sm^R$ | (1) |
| DH5α/λ*pir* | λ-pir cloning strain | (2) |
| TOP 10 F' | Cloning strain | Invitrogen |
| ***A. tumefaciens*** | | |
| C58 | Wild-type, nopaline-pTi, genomospecies G8 | (3) |
| C58-JX110 | ∆*crdS* ∆*celH-D* ∆*exoA* ∆*chvAB,* (CDGS-) | (4) |
| C58-JX137 | ∆*pruA* (ATU_RS05580, Atu1130) | (5) |
| C58-PMM29 | ∆*uppW* (ATU_RS11495, Atu2356) | (6) |
| C58-JX180 | ∆*uppY* (ATU_RS02370, Atu0481) | (6) |
| C58-JX185 | ∆*uppY* ∆*uppW* | (6) |
| C58-IPR12 | ∆*rfbA* (ATU_RS21635, Atu4615) | This study |
| C58-IPR19 | C58-JX180 (∆*uppY*), ∆*rfbA* | This study |
| C58-IPR6 | C58-PMM29 (∆*uppW*), ∆*rfbA* | This study |
| C58-IPR24 | C58-JX185 (∆*uppY* ∆*uppW*), ∆*rfbA* | This study |
| C58-IPR2 | ∆*rfbD* (ATU_RS21640, atu4616) | This study |
| C58-IPR8 | C58-JX180 (∆*uppY*), ∆*rfbD* | This study |
| C58-IPR21 | C58-PMM29 (∆*uppW*), ∆*rfbD* | This study |
| C58-IPR18 | C58-JX185 (∆*uppY* ∆*uppW*), ∆*rfbD* | This study |
| C58-IPR41 | ∆*rfbB* (ATU_RS21645, Atu4617) | This study |
| C58-IPR42 | ∆*rfbC* (ATU_RS21650, Atu4618) | This study |
| C58-IPR48 | ∆*exoC* (ATU_RS19035 Atu4074) | This study |
| C58-IPR49 | ∆*atu4608* (ATU_RS21610) | This study |
| C58-IPR43 | ∆*rfbB* ∆*rfbD* | This study |
| C58-IPR44 | ∆*rfbA* ∆*rfbD* | This study |
| C58-IPR45 | ∆*rfbC* ∆*rfbD* | This study |
| C58-IPR46 | ∆*rfbA,* ∆*rfbB,* ∆*rfbC,* ∆*rfbD* | This study |
| C58-IPR53 | ∆*exoC,* ∆*rfbD* | This study |
| C58-IPR54 | ∆*atu4608,* ∆*rfbD* | This study |
| C58-JX153 | JX110 (CDGS-), ∆*pruA* | (6) |
| C58-MCO070 | C58-JX151 (CDGS-), ∆*pruA*, ∆*uppY* | (6) |
| C58-MCO071 | C58-JX151 (CDGS-), ∆*pruA*, ∆*uppW* | (6) |
| C58-MCO072 | C58-JX151 (CDGS-), ∆*pruA*, ∆*uppY*, ∆*uppW* | (6) |
| C58-IPR14 | C58-JX151 (CDGS-), ∆*pruA,* ∆*rfbA* | This study |
| C58-IPR9 | C58-JX151 (CDGS-), ∆*pruA*, ∆*uppY,* ∆*rfbA* | This study |
| C58-IPR11 | C58-JX151 (CDGS-), ∆*pruA*, ∆*uppW,* ∆*rfbA* | This study |
| C58-IPR1 | C58-JX151 (CDGS-), ∆*pruA*, ∆*uppY*, ∆*uppW,* ∆*rfbA* | This study |
| C58-IPR20 | C58-JX151 (CDGS-), ∆*pruA,* ∆*rfbD* | This study |
| C58-IPR22 | C58-JX151 (CDGS-), ∆*pruA*, ∆*uppY,* ∆*rfbD* | This study |
| C58-IPR7 | C58-JX151 (CDGS-), ∆*pruA*, ∆*uppW,* ∆*rfbD* | This study |
| C58-IPR4 | C58-JX151 (CDGS-), ∆*pruA*, ∆*uppY*, ∆*uppW,* ∆*rfbD* | This study |
| C58-IPR36 | C58-JX151 (CDGS-), ∆*pruA,* ∆*rfbB* | This study |
| C58-IPR37 | C58-JX151 (CDGS-), ∆*pruA,* ∆*rfbC* | This study |
| C58-IPR50 | C58-JX151 (CDGS-), ∆*pruA,* ∆*exoC* | This study |

| | | |
|---|---|---|
| C58-IPR51 | C58-JX151 (CDGS-), Δ*pruA*, Δ*atu4608* | This study |
| C58-IPR38 | C58-JX151 (CDGS-), Δ*pruA*, Δ*rfbB* Δ*rfbD* | This study |
| C58-IPR39 | C58-JX151 (CDGS-), Δ*pruA*, Δ*rfbA* Δ*rfbD* | This study |
| C58-IPR40 | C58-JX151 (CDGS-), Δ*pruA*, Δ*rfbC* Δ*rfbD* | This study |
| C58-IPR47 | C58-JX151 (CDGS-), Δ*pruA*, Δ*rfbA*, Δ*rfbB*, Δ*rfbC*, Δ*rfbD* | This study |
| C58-IPR52 | C58-JX151 (CDGS-), Δ*pruA*, Δ*rfbD*, Δ*exoC* | This study |
| C58-IPR55 | C58-JX151 (CDGS-), Δ*pruA*, Δ*rfbD*, Δ*atu4608* | This study |
| C58-IPR83 | C58-JX151 (CDGS-), Δ*pruA*, Δ*rfbD*, Δ*flgE* | This study |
| C58-IPR86 | Δ*exoN* Δ*rfbD* | This study |
| C58-IPR87 | Δ*exoN* | This study |
| **Plasmids** | | |
| pNPTS138 | ColE1 suicide plasmid; *sacB*; $Km^R$ | (7) |
| pFD1 | Mariner *Himar1* suicide delivery plasmid, $Km^R$, $Ap^R$ | (8) |
| pSRKGm | Broad host range expression $P_{lac}$ expression vector plasmid, $lacI^Q$, $Gm^R$ | (9) |
| pBCH111 | pRA301::$P_{aopB}$-*lacZ*, $Spec^R$ | (10) |
| pBM205 | pBM205 [pSRKKm::com (FlaAT213C +FlaA's native promoter)] $Km^R$ | (11) |
| pBCH181 | pRA301::$P_{chvI}$-*lacZ*, $Spec^R$ | (10) |
| pPM107 | pKNG101::*flgE* knock-out fragment, upstream 500 bp and downstream 500 bp of genes, $Sm^R$ | (12) |
| pIR02 | pNPTS138; *rfbD* knock-out fragment, upstream 500bp and downstream 500bp of gene, $Km^R$ | This study |
| pIR03 | pNPTS138; *rfbA* knock-out fragment, upstream 500bp and downstream 500bp of gene, $Km^R$ | This study |
| pIR10 | pNPTS138; *rfbC* knock-out fragment, upstream 500bp and downstream 500bp of gene, $Km^R$ | This study |
| pIR11 | pNPTS138; *rfbB rfbD* knock-out fragment, upstream 500bp and downstream 500bp of genes, $Km^R$ | This study |
| pIR12 | pNPTS138; *rfbA rfbD* knock-out fragment, upstream 500bp and downstream 500bp of genes, $Km^R$ | This study |
| pIR13 | pNPTS138; *rfbA rfbB rfbC rfbD* knock-out fragment, upstream 500bp and downstream 500bp of genes, $Km^R$ | This study |
| pIR14 | pNPTS138; *rfbB* knock-out fragment, upstream 500bp and downstream 500bp of gene, $Km^R$ | This study |
| pIR15 | pNPTS138; *exoC* knock-out fragment, upstream 500bp and downstream 500bp of genes, $Km^R$ | This study |
| pIR16 | pNPTS138; *atu4608* knock-out fragment, upstream 500bp and downstream 500bp of genes, $Km^R$ | This study |
| pIR72 | pNPTS138; *exoN* knock-out fragment, upstream 500bp and downstream 500bp of genes, $Km^R$ | This study |
| pIR04 | pSRKGm carrying $P_{lac}$-*rfbA*, $Gm^R$ | This study |
| pIR06 | pSRKGm carrying $P_{lac}$-*rfbD*, $Gm^R$ | This study |
| pIR07 | pSRKGm carrying $P_{lac}$-*motAB*, $Gm^R$ | This study |

| | | |
|---|---|---|
| pIR09 | pSRKGm carrying $P_{lac}$-*atu4608*, $Gm^R$ | This study |
| pIR18 | pSRKGm carrying $P_{lac}$-*rfbB*, $Gm^R$ | This study |
| pIR19 | pSRKGm carrying $P_{lac}$-*rfbC*, $Gm^R$ | This study |
| pIR21 | pSRKGm carrying $P_{lac}$-*rfbCBDA*, $Gm^R$ | This study |
| pIR73 | pSRKGm carrying $P_{lac}$-*rfbCBA*, $Gm^R$ | This study |
| pIR74 | pSRKGm carrying $P_{lac}$-*rfbDA*, $Gm^R$ | This study |

**Table S3 Oligonucleotides used in study**

| Oligos | Sequence | Use |
|---|---|---|
| IR131 / pNPTS_FC .FOR | gcatgcgtcgaccctctagtcaagg | pNPTS138 derived knock-out constructs |
| IR132 / pNPTS_FC .REV | actagtgagtcgtattacgtagcttggcg | pNPTS138 derived knock-out constructs |
| IR098/ pSRK_FC. FOR | actagtggatcccccgggctgc | pSRKGm derived complementation constructs |
| IR099/ pSRK_FC. REV | catatgctgtttcctgtgtgaaattgttatcc | pSRKGm derived complementation constructs |
| JEH89 | gcgttggccgattcattaatgca | sequencing |
| JEH90 | gtcaattattacctccacgggga | sequencing |
| M13_F | cgccagggttttcccagtcacgac | sequencing |
| M13_R | tcacacaggaaacagctatgac | sequencing |
| IPCR A F | cgggtatcgctcttgaaggga | sequencing |
| IPCR A R | ctcgaattgacgcgtcgag | sequencing |
| IR104 | aagctacgtaatacgactcactagtattccacttcccggaaaagc t | pIR02, pIR12 |
| IR105 | cctttttggttgccagccgcatacgat | pIR02 |
| IR106 | gcggctggcaaccaaaaaggaaaagtcatgaaggg | pIR02 |
| IR107 | gactagagggtcgacgcatgccggtttttcttcgatggaaagagc ac | pIR02 , pIR11 |
| IR108 | aagctacgtaatacgactcactagtgtttcggaggcaaccgaca at | pIR03 |
| IR109 | aaggtgcggcgatgcccttcatgacttttcctttttgg | pIR03 |
| IR110 | gaagggcatcgccgcaccttaaaggatgg | pIR03 |
| IR111 | gactagagggtcgacgcatgcgcggggcaaaagccatga | pIR03, pIR13, pIR12 |
| IR114 | ttcacacaggaaacagcatatgaagggcatcattctggccggc | pIR04 |
| IR115 | cagcccgggggatccactagtttaaggtgcggcaagctttcttaa ataaacgc | pIR04, pIR74 |
| IR116 | cacacaggaaacagcatatggtatgcggctggcagttaccggc a | pIR06, pIR74 |
| IR117 | agcccgggggatccactagttcatgacttttcctttttggtctgcgca agagc | pIR06 |
| IR120 | cacacaggaaacagcatatgaatattgtaattggacttataatca ccttcggctgca | pIR07 |
| IR121 | tcaccttgcgtcatgccgccttgttttcgccgc | pIR07 |
| IR122 | gcggcatgacgcaaggtgatgaaatgagtgaaggcg | pIR07 |
| IR123 | gcagcccgggggatccactagttcacccccgatcagcctcaag cag | pIR07 |
| IR129 | cacacaggaaacagcatatgttgaacccagatggacactatga tcatgc | pIR09 |
| IR130 | agcccgggggatccactagtttatttccgccgatcaccgtggaaa atc | pIR09 |
| IR133 | cgtaatacgactcactagtagggccaggaatggcgttcc | pIR10, pIR13 |

| | | |
|---|---|---|
| IR134 | tcatcagttctcttcgcggatcctgtctcatctgcc | pIR10 |
| IR135 | gacaggatccgcgaagagaactgatgatgcgcgttctagttacc | pIR10 |
| IR136 | actagagggtcgacgcatgcggctttcgaggcggaataggg | pIR10 |
| IR137 | gtaatacgactcactagtgttttcaaacgagagtggtttcgcgaaaatgtag | pIR11, pIR14 |
| IR138 | ttttccttttggttagaacgcgcatcatcagttctcggt | pIR11 |
| IR139 | gatgcgcgttctaaccaaaaaggaaaagtcatgaagggcatcatt | pIR11 |
| IR140 | ccctttgtcagtagaacgcgcatcatcagttctcggtgt | pIR14 |
| IR141 | tgcgcgttctactgacaaagggctaatcgtatgcggc | pIR14 |
| IR142 | tagagggtcgacgcatgccacgaagttattgccatattccgagtagac | pIR14 |
| IR143 | aggtgcggcttgcacttcgcggatcctgtctcat | pIR13 |
| IR144 | gcgaagtgcaagccgcaccttaaaggatgggc | pIR13 |
| IR145 | ggtgcggctgccagccgcatacgattagcc | pIR12 |
| IR146 | cggctggcagccgcaccttaaaggatggg | pIR12 |
| IR147 | gtaatacgactcactagtaaattcgactggcagcgtccaaagc | pIR16 |
| IR148 | ctctgatgttgaacccagatcggcggaaataatgagcgtggt | pIR16 |
| IR149 | gccgatctgggttcaacatcagagtacaaacctatgtttggtataaaaatgac | pIR16 |
| IR150 | tagagggtcgacgcatgcatcgagcatcccaaggcctacg | pIR16 |
| IR151 | cacaggaaacagcatatgcaactagaagtatttccgatatcaggc | pIR19 |
| IR152 | cccgggggatccactagttcagttctcggtgtttcggtaggtg | pIR19 |
| IR153 | cacaggaaacagcatatgcgcgttctagttaccggcg | pIR18 |
| IR154 | cccgggggatccactagtttagccctttgtcagaacgccaagg | pIR18 |
| IR157 | gtaatacgactcactagtatgcgcggaaaggatcgcct | pIR15 |
| IR158 | tgaaccaatgatcaagactactgtcattacctgataacgtccgccc | pIR15 |
| IR159 | cgttatcaggtaatgacagtagtcttgatcattggttcaatggcctttcg | pIR15 |
| IR160 | tagagggtcgacgcatgccggccgcctggtcgtgc | pIR15 |
| IR263 | cagccgcatacgattagccctttg | pIR73 |
| IR264 | cagaccaaaaaggaaaagtcatgaagggcatcattctgg | pIR73 |
| IR259 | GTAATACGACTCACTAGTtggcgacatcggcggattgac | pIR72 |
| IR260 | cgatggacctggtccaggcggcctgacgttcatcg | pIR72 |
| IR261 | ccgcctggaccaggtccatcgtgttctccttcatcaagg | pIR72 |
| IR262 | TAGAGGGTCGACGCATGCcgcatcggcgaggattatatcctgct | pIR72 |

**Supplementary Methods**

**Plasmid construction**

pIR02 [pNPTS138 Δ*rfbD* (ATU_RS21640/atu4616) (*aph*I $Km^R$) ] was constructed by performing Gibson assembly (13) using a 500 bp product amplified from the region upstream of *rfbD* using oligos IR104 and IR105 on C58 gDNA, with a 500 bp product amplified from the region downstream of *rfbD* using oligos IR106 and IR107 on C58 gDNA, and with pNPTS138 digested with *Nde*I and *Spe*I.

pIR03 [pNPTS138 Δ*rfbA* (ATU_RS21635/atu4615) (*aph*I $Km^R$)] was constructed by performing Gibson assembly using the 50 0bp amplified product from upstream region of *rfbA* using oligos IR108 and IR109 on C58 gDNA, with a 500 bp product amplified from the region downstream of *rfbA* using oligos IR110 and IR111 on C58 gDNA, along with pNPTS138 digested with *Nde*I and *Spe*I.

pIR10 [pNPTS138 Δ*rfbC* (ATU_RS21650/atu4618) (*aph*I $Km^R$) ] was constructed by first amplifying the 500 bp product from the upstream region of *rfbC* using oligos IR133 and IR134 on C58 gDNA and a 500bp product amplified from the region downstream of *rfbC* using oligos IR135 and IR136 on C58 gDNA. SOE PCR was performed with the upstream and downstream products using oligos IR133 and IR136. FastCloning (14) was then performed using the 1 kb *rfbC* knock-out fragment and the PCR product of pNPTS138 amplified using oligos pNPTS138_FC.FOR and pNPTS138_FC.REV.

pIR11 [pNPTS138 Δ*rfbB* D*rfbD* (ATU_RS21645/atu4617, ATU_RS21640/atu4616, respectively) (*aph*I $Km^R$) ] was constructed by first amplifying the 500 bp product from the upstream region of *rfbB/D* using oligos IR137 and IR138 and a 500 bp product amplified from the region downstream of *rfbB/D* using oligos IR139 and IR107, each using C58 gDNA as the template. SOE PCR was performed with the upstream and downstream products using oligos IR137 and IR107. FastCloning was then performed using the 1kb *rfbB/D* knock-out fragment and the PCR product of pNPTS138 amplified using oligos pNPTS138_FC.FOR and pNPTS138_FC.REV.

pIR12 [pNPTS138 Δ*rfbA* Δ*rfbD* (ATU_RS21635/atu4615, ATU_RS21640/atu4616, respectively) (*aph*I $Km^R$) ] was constructed by first amplifying the 500 bp product from the upstream region of *rfbD/A* using oligos IR104 and IR145 and a 500 bp product amplified from the region downstream of *rfbD/A* using oligos IR111 and IR146, each using C58 gDNA as the template . SOE PCR was performed with the upstream and downstream products using oligos IR104 and IR111. FastCloning was then performed using the 1 kb *rfbD/A* knock-out fragment and the PCR product of pNPTS138 amplified using oligos pNPTS138_FC.FOR and pNPTS138_FC.REV.

pIR13 [pNPTS138 Δ*rfbCBDA* (ATU_RS21635/atu4615 through ATU_RS21650/atu4618) (*aph*I $Km^R$) ] was constructed by first amplifying the 500 bp product from the upstream region of *rfbCBDA* using oligos IR133 and IR143 and a 500 bp product amplified from the region downstream of *rfbCBDA* using oligos IR111 and IR144, each using C58 gDNA as the template . SOE PCR was performed with the upstream and downstream products using oligos IR133 and IR111. FastCloning was then performed using the 1 kb *rfbCBDA* knock-out fragment and the PCR product of pNPTS138 amplified using oligos pNPTS138_FC.FOR and pNPTS138_FC.REV.

pIR14 [pNPTS138 Δ*rfbB* (ATU_RS21645/atu4617) (*aph*I $Km^R$) ] was constructed by amplifying the 500 bp product from the upstream region of *rfbB* using oligos IR137 and IR140 and a 500 bp product amplified from the region downstream of *rfbB* using oligos IR141 and IR142, each using C58 gDNA as the template . SOE PCR was performed with the upstream and downstream products using oligos IR137 and IR142. FastCloning was then performed using the 1 kb *rfbB* knock-out fragment and the PCR product of pNPTS138 amplified using oligos pNPTS138_FC.FOR and pNPTS138_FC.REV.

pIR15 [pNPTS138 Δ*exoC* (ATU_RS19035 Atu4074) (*aph*I $Km^R$) ] was constructed by amplifying the 500 bp product from the upstream region of *exoC* using oligos IR157 and IR158 and a 500 bp product amplified from the region downstream of *exoC* using oligos IR159 and IR160, each using C58 gDNA as the template . SOE PCR was performed with the upstream and downstream products using oligos IR157 and IR160. FastCloning was then performed using the 1 kb *exoC* knock-out fragment and the PCR product of pNPTS138 amplified using oligos pNPTS138_FC.FOR and pNPTS138_FC.REV.

pIR16 [pNPTS138 Δ*atu4608* (ATU_RS21610/Atu4608) (*aph*I $Km^R$) ] was constructed by amplifying the 500 bp product from the upstream region of *atu4608* using oligos IR147 and IR148 and a 500 bp product amplified from the region downstream of *atu4608* using oligos IR149 and IR150, each using C58 gDNA as the template . SOE PCR was performed with the upstream and downstream products using oligos IR147 and IR150. FastCloning was then performed using the 1 kb *atu4608* knock-out fragment and the PCR product of pNPTS138 amplified using oligos pNPTS138_FC.FOR and pNPTS138_FC.REV.

pIR04 [pSRKGm $P_{lac}$*::rfbA (*ATU_RS21635/atu4615) $Gm^R$] was made by amplifying *rfbA* from C58 gDNA using oligos IR114 and IR115, amplifying the pSRKGm template with oligos pSRKGm_FC.FOR and pSRKGm_FC.REV, and performing Fast Cloning with the PCR products.

pIR06 [pSRKGm $P_{lac}$*::rfbD (*ATU_RS21640/atu4616) $Gm^R$ ] was made by amplifying *rfbA* from C58 gDNA using oligos IR116 and IR117, amplifying the pSRKGm template with oligos pSRKGm_FC.FOR and pSRKGm_FC.REV, and performing Fast Cloning with the PCR products.

pIR09 [pSRKGm $P_{lac}$*::atu4608 (*ATU_RS21610/Atu4608) $Gm^R$ ] was made by amplifying *atu4608* from C58 gDNA using oligos IR129 and IR130, amplifying the pSRKGm template with oligos pSRKGm_FC.FOR and pSRKGm_FC.REV, and performing Fast Cloning with the PCR products.

pIR18 [pSRKGm $P_{lac}$::*rfbB (*ATU_RS21645/atu4617) Gm$^R$ ] was made by amplifying *rfbB* from C58 gDNA using oligos IR153 and IR154, amplifying the pSRKGm template with oligos pSRKGm_FC.FOR and pSRKGm_FC.REV, and performing Fast Cloning with the PCR products.

pIR19 [pSRKGm $P_{lac}$::*rfbC* (ATU_RS21650/atu4618) Gm$^R$ ] was made by amplifying *rfbC* from C58 gDNA using oligos IR151 and IR152, amplifying the pSRKGm template with oligos pSRKGm_FC.FOR and pSRKGm_FC.REV, and performing Fast Cloning with the PCR products.

pIR21 [pSRKGm $P_{lac}$::*rfbCBDA (*ATU_RS21635/atu4615 through ATU_RS21650/atu4618) Gm$^R$] was made by amplifying *rfbCBDA* from C58 gDNA using oligos IR153 and IR115, amplifying the pSRKGm template with oligos pSRKGm_FC.FOR and pSRKGm_FC.REV, and performing Fast Cloning with the PCR products.

pIR74 [pSRKGm carrying $P_{lac}$-*rfbDA*, Gm$^R$ (ATU_RS21640/atu4616 and ATU_RS21650/atu4618)] was made by amplifying *rfbDA* fragment form C58 gDNA using oligos IR115 and IR116, amplifying the pSRKGm template with oligos pSRKGm_FC.FOR and pSRKGm_FC.REV, and performing Fast Cloning with the PCR products.

pIR73 [pSRKGm carrying $P_{lac}$-*rfbCBA*, Gm$^R$  (ATU_RS21635/atu4615 through ATU_RS21650/atu4618)] was made by site-directed mutagenesis of the pIR21 vector using oligos IR263 and IR264 to exclude the *rfbD* gene and re-ligate the plasmid back together using the NEB Q5 site-directed mutagenesis kit.

pIR72 [pNPTS138; *exoN* knock-out fragment, upstream 500bp and downstream 500bp of genes (*aph*I Km$^R$)(ATU_RS18920/atu4050)] was made by amplifying 500bp upstream and downstream fragments from C58 gDNA using oligo pairs of IR259 + IR260 and IR261 and IR262. These fragments were then fused using SOE PCR with oligos IR259 and IR262. FastCloning was then performed using the 1kb *exoN* knock-out fragment and the PCR product of pNPTS138 amplified using oligos pNPTS138_FC.FOR and pNPTS138_FC.REV.

**Strain Construction**

C58 CDGS- Δ*pruA*Δ*uppY*Δ*uppW*Δ*rfbA* (IPR1) was generated by in frame markerless deletion (7), using S17-1 to conjugate pIR03 [pNPTS138 Δ*rfbA* (ATU_RS21635/atu4615) (Km$^{R}$)] into C58 CDGS- Δ*pruA* Δ*uppY* Δ*uppW*.

C58 CDGS- Δ*pruA* Δ*rfbA* (IPR14) was generated by in frame markerless deletion, using S17-1 to conjugate pIR03 [pNPTS138 Δ*rfbA* (ATU_RS21635/atu4615) (Km$^{R}$)] into C58 CDGS- Δ*pruA*.

C58 CDGS- Δ*pruA*Δ*uppY*Δ*rfbA* (IPR9) was generated by in frame markerless deletion, using S17-1 to conjugate pIR03 [pNPTS138 Δ*rfbA* (ATU_RS21635/atu4615) (Km$^{R}$)] into C58 CDGS- Δ*pruA* Δ*uppY*.

C58 CDGS- Δ*pruA*Δ*uppW*Δ*rfbA* (IPR11) was generated by in frame markerless deletion, using S17-1 to conjugate pIR03 [pNPTS138 Δ*rfbA* (ATU_RS21635/atu4615) (Km$^{R}$)] into C58 CDGS- Δ*pruA*Δ*uppW*.

C58 Δ*rfbA* (IPR12) was generated by in frame markerless deletion, using S17-1 to conjugate pIR03 [pNPTS138 Δ*rfbA* (ATU_RS21635/atu4615) (Km$^{R}$)] into C58.

C58 Δ*uppY*Δ*rfbA* (IPR19) was generated by in frame markerless deletion, using S17-1 to conjugate pIR03 [pNPTS138 Δ*rfbA* (ATU_RS21635/atu4615) (Km$^{R}$)] into C58 Δ*uppY*.

C58 Δ*uppW*Δ*rfbA* (IPR6) was generated by in frame markerless deletion, using S17-1 to conjugate pIR03 [pNPTS138 Δ*rfbA* (ATU_RS21635/atu4615) (Km$^{R}$)] into C58 Δ*uppW*.

C58 Δ*uppY*Δ*uppW*Δ*rfbA* (IPR24) was generated by in frame markerless deletion, using S17-1 to conjugate pIR03 [pNPTS138 Δ*rfbA* (ATU_RS21635/atu4615) (Km$^{R}$)] into C58 Δ*uppY*Δ*uppW*.

C58 Δ*rfbD* (IPR12) was generated by in frame markerless deletion, using S17-1 to conjugate pIR02 [pNPTS138 Δ*rfbD* (ATU_RS21640/atu4616) (Km$^{R}$)] into C58.

C58 Δ*uppY*Δ*rfbD* (IPR8) was generated by in frame markerless deletion, using S17-1 to conjugate pIR02 [pNPTS138 Δ*rfbD* (ATU_RS21640/atu4616) ($Km^R$)] into C58 Δ*uppY*.

C58 Δ*uppW* Δ*rfbD* (IPR21) was generated by in frame markerless deletion, using S17-1 to conjugate pIR02 [pNPTS138 Δ*rfbD* (ATU_RS21640/atu4616) ($Km^R$)] into C58 Δ*uppW*.

C58 Δ*uppY* Δ*uppW* Δ*rfbD* (IPR18) was generated by in frame markerless deletion, using S17-1 to conjugate pIR02 [pNPTS138 Δ*rfbD* (ATU_RS21640/atu4616) ($Km^R$)] into C58 Δ*uppY* Δ*uppW*.

C58 CDGS- Δ*pruA*Δ*rfbD* (IPR20) was generated by in frame markerless deletion, using S17-1 to conjugate pIR02 [pNPTS138 Δ*rfbD* (ATU_RS21640/atu4616) ($Km^R$)] into C58 CDGS- Δ*pruA*.

C58 CDGS- Δ*pruA*Δ*uppY*Δ*rfbD* (IPR22) was generated by in frame markerless deletion, using S17-1 to conjugate pIR02 [pNPTS138 Δ*rfbD* (ATU_RS21640/atu4616) ($Km^R$)] into C58 CDGS- Δ*pruA* Δ*uppY*.

C58 CDGS- Δ*pruA*Δ*uppW*Δ*rfbD* (IPR7) was generated by in frame markerless deletion, using S17-1 to conjugate pIR02 [pNPTS138 Δ*rfbD* (ATU_RS21640/atu4616) ($Km^R$)] into C58 CDGS- Δ*pruA*Δ*uppW*.

C58 CDGS- Δ*pruA*Δ*uppY*Δ*uppW*Δ*rfbD* (IPR4) was generated by in frame markerless deletion, using S17-1 to conjugate pIR02 [pNPTS138 Δ*rfbD* (ATU_RS21640/atu4616) ($Km^R$] into C58 CDGS- Δ*pruA*Δ*uppY*Δ*uppW*.

C58 CDGS- Δ*pruA*Δ*rfbB* (IPR36) was generated by in frame markerless deletion, using S17-1 to conjugate pIR14 [pNPTS138 Δ*rfbB* (ATU_RS21645/atu4617) ($Km^R$)] into C58 CDGS- Δ*pruA*.

C58 CDGS- Δ*pruA*Δ*rfbC* (IPR37) was generated by in frame markerless deletion, using S17-1 to conjugate pIR10 [pNPTS138 D*rfbC* (ATU_RS21650/atu4618) ($Km^R$)] into C58 CDGS- Δ*pruA*.

C58 CDGS- Δ*pruA*Δ*rfbB*Δ*rfbD* (IPR38) was generated by in frame markerless deletion, using S17-1 to conjugate pIR11 [pNPTS138 Δ*rfbB* Δ*rfbD* (ATU_RS21645/atu4617, ATU_RS21640/atu4616, respectively) ($Km^R$)] into C58 CDGS- Δ*pruA*.

C58 CDGS- Δ*pruA*Δ*rfbA*Δ*rfbD* (IPR39) was generated by in frame markerless deletion, using S17-1 to conjugate pIR12 [pNPTS138 Δ*rfbA* Δ*rfbD* (ATU_RS21635/atu4615, ATU_RS21640/atu4616, respectively) ($Km^R$)] into C58 CDGS- Δ*pruA*.

C58 CDGS- Δ*pruA*Δ*rfbC*Δ*rfbD* (IPR40) was generated by in frame markerless deletion, using S17-1 to conjugate pIR10 [pNPTS138 Δ*rfbC* (ATU_RS21650/atu4618) ($Km^R$)] into C58 CDGS- Δ*pruA*Δ*rfbD*.

C58 Δ*rfbB* (IPR41) was generated by in frame markerless deletion, using S17-1 to conjugate pIR14 [pNPTS138 Δ*rfbB* (ATU_RS21645/atu4617) ($Km^R$)] into C58.

C58 Δ*rfbC* (IPR42) was generated by in frame markerless deletion, using S17-1 to conjugate pIR10 [pNPTS138 Δ*rfbC* (ATU_RS21650/atu4618) ($Km^R$)] into C58.

C58 Δ*rfbB*Δ*rfbD* (IPR43) was generated by in frame markerless deletion, using S17-1 to conjugate pIR11 [pNPTS138 Δ*rfbB* Δ*rfbD* (ATU_RS21645/atu4617, ATU_RS21640/atu4616, respectively) ($Km^R$)] into C58.

C58 Δ*rfbA*Δ*rfbD* (IPR44) was generated by in frame markerless deletion, using S17-1 to conjugate pIR12 [pNPTS138 Δ*rfbA*Δ*rfbD* (ATU_RS21635/atu4615, ATU_RS21640/atu4616, respectively) ($Km^R$)] into C58.

C58 Δ*rfbC*Δ*rfbD* (IPR45) was generated by in frame markerless deletion, using S17-1 to conjugate pIR10 [pNPTS138 Δ*rfbC* (ATU_RS21650/atu4618) ($Km^R$)] into C58 Δ*rfbD*.

C58 Δ*rfbCBDA* (IPR46) was generated by in frame markerless deletion, using S17-1 to conjugate pIR13 [pNPTS138 Δ*rfbCBDA* (ATU_RS21635/atu4615 through ATU_RS21650/atu4618) ($Km^R$)] into C58.

C58 CDGS- Δ*pruA*Δ*rfbCBDA* (IPR47) was generated by in frame markerless deletion, using S17-1 to conjugate pIR13 [pNPTS138 Δ*rfbCBDA* (ATU_RS21635/atu4615 through ATU_RS21650/atu4618) ($Km^R$)] into C58 CDGS- Δ*pruA*.

C58 Δ*exoC* (IPR48) was generated by in frame markerless deletion, using S17-1 to conjugate pIR15 [pNPTS138 Δ*exoC* (ATU_RS19035 Atu4074) ($Km^R$)] into C58.

C58 Δ*atu4608* (IPR49) was generated by in frame markerless deletion, using S17-1 to conjugate pIR16 [pNPTS138 Δ*atu4608* (ATU_RS21610/Atu4608) ($Km^R$)] into C58.

C58 CDGS- Δ*pruA*Δ*exoC* (IPR50) was generated by in frame markerless deletion, using S17-1 to conjugate pIR15 [pNPTS138 Δ*exoC* (ATU_RS19035 Atu4074) ($Km^R$)] into C58 CDGS- Δ*pruA*.

C58 CDGS- Δ*pruA*Δ*atu4608* (IPR51) was generated by in frame markerless deletion, using S17-1 to conjugate pIR16 [pNPTS138 Δ*atu4608* (ATU_RS21610/Atu4608) ($Km^R$)] into C58 CDGS- Δ*pruA*.

C58 CDGS- Δ*pruA* Δ*exoC* Δ*rfbD* (IPR52) was generated by in frame markerless deletion, using S17-1 to conjugate pIR15 [pNPTS138 Δ*exoC* (ATU_RS19035 Atu4074) ($Km^R$)] into C58 CDGS- Δ*pruA*Δ*rfbD*.

C58 Δ*exoC*Δ*rfbD* (IPR53) was generated by in frame markerless deletion, using S17-1 to conjugate pIR15 [pNPTS138 Δ*exoC* (ATU_RS19035 Atu4074) ($Km^R$)] into C58 Δ*rfbD*.

C58 Δ*atu4608*Δ*rfbD* (IPR54) was generated by in frame markerless deletion, using S17-1 to conjugate pIR16 [pNPTS138 Δ*atu4608* (ATU_RS21610/Atu4608) ($Km^R$)] into C58 Δ*rfbD*.

C58 CDGS- Δ*pruA*Δ*atu4608*Δ*rfbD* (IPR55) was generated by in frame markerless deletion, using S17-1 to conjugate pIR16 [pNPTS138 Δ*atu4608* (ATU_RS21610/Atu4608) ($Km^R$)] into C58 CDGS- Δ*pruA*Δ*rfbD*.

C58 CDGS- Δ*pruA*Δ*rfbD*Δ*flgE* (IPR83) was generated by in-frame markerless deletion, using S17-1 to conjugate pPM107 [pKNG101::*flgE* knock-out fragment, upstream 500 bp and downstream 500 bp of genes, $Sm^R$] into C58 CDGS- Δ*pruA*Δ*rfbD*.

C58 Δ*exoN*Δ*rfbD* (IPR86) was generated by in-frame markerless deletion, using S17-1 to conjugate pIR72 [pNPTS138; *exoN* knock-out fragment, upstream 500 bp and downstream 500 bp of genes, $Km^R$] into C58 Δ*rfbD*.

C58 Δ*exoN* (IPR87) was generated by in-frame markerless deletion, using S17-1 to conjugate pIR72 [pNPTS138; *exoN* knock-out fragment, upstream 500 bp and downstream 500 bp of genes, $Km^R$] into C58.
